# Single cell genome and epigenome co-profiling reveals hardwiring and plasticity in breast cancer

**DOI:** 10.1101/2024.09.06.611519

**Authors:** Kaile Wang, Yun Yan, Heba Elgamal, Jianzhuo Li, Chenling Tang, Shanshan Bai, Zhenna Xiao, Emi Sei, Yiyun Lin, Junke Wang, Jessica Montalvan, Changandeep Nagi, Alastair M. Thompson, Nicholas Navin

## Abstract

Understanding the impact of genetic alterations on epigenomic phenotypes during breast cancer progression is challenging with unimodal measurements. Here, we report wellDA-seq, the first high-genomic resolution, high-throughput method that can simultaneously measure the whole genome and chromatin accessibility profiles of thousands of single cells. Using wellDA-seq, we profiled 22,123 single cells from 2 normal and 9 tumors breast tissues. By directly mapping the epigenomic phenotypes to genetic lineages across cancer subclones, we found evidence of both genetic hardwiring and epigenetic plasticity. In 6 estrogen-receptor positive breast cancers, we directly identified the ancestral cancer cells, and found that their epithelial cell-of-origin was Luminal Hormone Responsive cells. We also identified cell types with copy number aberrations (CNA) in normal breast tissues and discovered non-epithelial cell types in the microenvironment with CNAs in breast cancers. These data provide insights into the complex relationship between genetic alterations and epigenomic phenotypes during breast tumor evolution.

## Introduction

Breast cancer is the most frequently diagnosed cancer in women worldwide^1^ and has been shown to exhibit extensive intratumoral genetic heterogeneity (ITH) and epigenetic and transcriptional diversity within the cancer cells^2,3^. However, how the genetic alterations manifest in phenotypic changes that are conferred through epigenetic and transcriptional programs remains a major unresolved question in the breast cancer field. A fundamental question is how specific genomic alterations acquiring in cancer subclones remodel the chromatin landscape^4^. Using bulk profiling methods, previous pan-cancer studies have shown that different cancer types with specific genetic backgrounds have different chromatin accessibility profiles^5,6^. However, ITH^7^ presents a major technical challenge for bulk genomic analysis of human tumors, when quantifying the impact of genomic changes on epigenetic regulation, which is critical for understanding tumor initiation and progression. Another key question is whether clonal lineages with specific genetic alterations ‘hardwire’ cancer cells towards a specific set of tumorigenic phenotypes, or alternatively, whether the cancer cells also exhibit phenotypic plasticity that is independent of the genetic lineages. Delineating these relationships requires directly measuring the genotype and phenotype (e.g., epigenome) of a single cell at scale to resolve these complex associations.

Breast cancer is commonly classified into estrogen receptor positive (ER+), progesterone receptor positive (PR+), human epidermal growth factor receptor positive (HER2+), and triple-negative breast cancer (TNBC)^8^. Normal breast tissue is composed of three different epithelial cells including basal myoepithelial cells (Basal), luminal hormone-responsive epithelial cells (LumHR) and luminal secretory epithelial cells (LumSec)^9,10^. However determining the cell-of-origin for each breast cancer subtype has remained challenging using canonical marker genes, since cancer cells can show plasticity and potentially switch lineages during cancer progression^11–13^. While high-throughput single cell DNA sequencing (scDNA-seq) methods can resolve clonal lineages in cancers and identify early ancestral populations, they cannot provide phenotypic information on the subpopulations that are ordered in these lineages^14–16^. Conversely, while high-throughput single cell ATAC sequencing (scATAC-seq) methods can provide phenotypic information and cell type identities, they cannot provide accurate genotypic information^17–20^.

To simultaneously capture the genome and epigenome information of single cells, computational tools have been developed to infer DNA copy number information from scATAC-seq^19,21–23^. While seemingly practical, these methods are limited to resolve larger chromosome events and have many false-positive CNA events due to the uneven and sparse ATAC read distribution across the genome. More recently the development of scGET-seq^24^ and dTag^25^ have improved the genomic resolution of whole genome copy number data by capturing additional regions from heterochromatin. However, a fundamental technical challenge persists: the need to remove nucleosome proteins to expose as many DNA regions as possible for DNA profiling, while simultaneously preserving cell or nuclear membrane proteins to maintain cell integrity for subsequent single-cell sequencing. This paradox has hindered methods for achieving high-genomic resolution CNA data to delineate the subclonal structure of tumors and study the interplay between genotypes and chromatin accessibility.

To overcome these challenges, we developed a nano**well D**NA&**A**TAC sequencing (wellDA-seq) method that can simultaneously measure genome-wide copy number profiles and chromatin accessibility from thousands of single cells in parallel. The compartmentalization of single cells in nanowells allows wellDA-seq to fully remove chromatin from DNA for high-resolution whole genome profiling, in addition to measuring chromatin accessibility. We applied wellDA-seq to ER+ breast cancers, which revealed both genetic hardwiring and phenotypic plasticity in subclones, the cells-of-origin of breast cancer cells originated from the LumHR lineages, and cis changes in chromatin accessibility that were associated with subclonal CNA events.

## Results

### Overview of the wellDA-seq method

wellDA-seq utilizes two distinct Tn5 transposomes (Tn5-1 and Tn5-2) with different adapter sets to specifically label the open chromatin regions (ATAC profile) and the remaining closed regions (DNA profile) of the entire genome (**Fig. 1a**). To perform wellDA-seq, the single cell suspensions are first permeabilized and then tagmented with the first Tn5 transposome (Tn5-1) to label the open chromatin regions (**Fig. 1a, step I**). After this tagmentation step, the cells are washed and stained with DAPI, and then loaded into a nanowell chip with 5,184 nanowells using a multi-channel micro-dispenser. Nanowells with single cells (1,200-2,600) are selected by imaging all wells to avoid doublets, degraded nuclei and empty wells (**Fig. 1a, step II**). Each single cell in the nanowell is then fully lysed with a protease-based buffer to completely remove the chromatin from the DNA. Next, a second set of the Tn5 transposome (Tn5-2) is used to perform the second round of tagmentation for labelling the previously closed chromatin regions across the genome. A set of 72 X 72 DNA index primers and 72 X 72 ATAC index primers matching with the transposome adaptors of both Tn5 transposomes, are added to the nanowells for subsequent PCR amplification reactions. By using unique combinations of DNA and ATAC indices, the molecules of the open and closed regions from the same cell are labelled with a unique cell barcode during PCR amplification (**Fig. 1a, step III**). The PCR products of each single cell from the entire chip are then pooled together, followed by a PCR amplification step to enrich the ATAC and DNA modalities with modality-specific primers, respectively. Finally, the enriched sequencing libraries are used for next-generation sequencing and downstream data analysis (**Fig. 1a, step IV – V, Extended Data Fig. 1**).

**Fig. 1.**
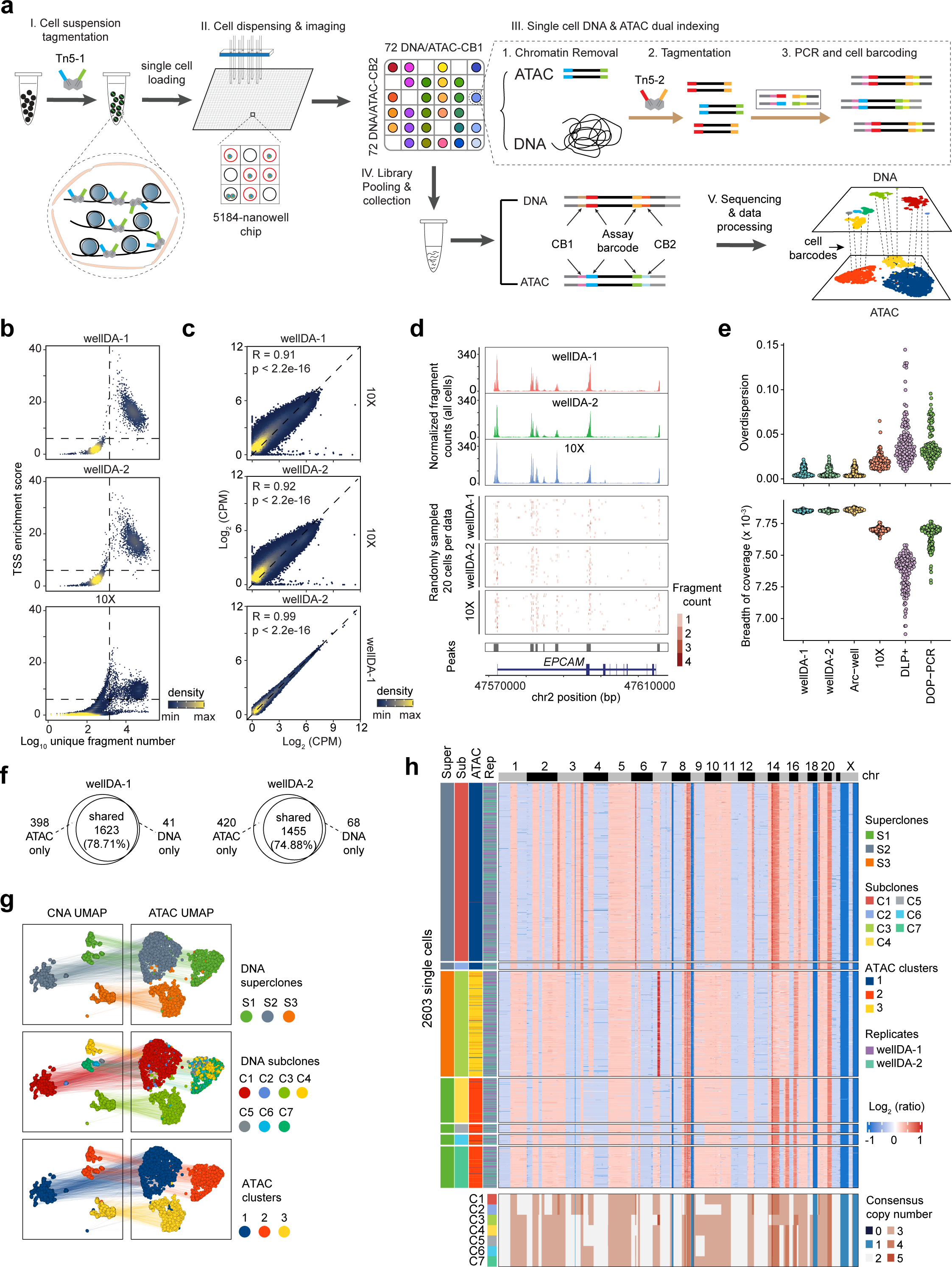
Workflow and technical performance of wellDA-seq. **a.** Experimental workflow of wellDA-seq using a nanowell microdispensing system. **b**. TSS enrichment score and unique fragment number of scATAC data from two wellDA-seq experiments (wellDA-1 and wellDA-2) and a 10X Genomics (10X) experiment of the MDA231 cell line. Each dot represents one nanowell or one microdroplet. **c.** Pearson correlation coefficient of the aggregated counts per million (CPM) fragments within peaks for the MDA231 cell line profiled by wellDA-seq and 10X. **d.** Comparison of scATAC profiles from the wellDA-seq and 10X in the *EPCAM* gene region of chromosome 2. The upper panels show the aggregated profiles of all cells, and the lower panels show fragments present in 20 random cells of each experiment. **e.** Comparison of the overdispersion metrics for the genomic bin counts and the breadth of coverage metrics of the scDNA data from wellDA-seq and four other unimodal scDNA-seq methods. **f.** Venn diagrams showing the number of cells with DNA and/or ATAC data that were profiled by the two wellDA-seq experiments. **g.** UMAP of the CNA modality (left panel) and ATAC modality (right panel) colored by the DNA superclones (top panel), the DNA subclones (middle panel), and the ATAC clusters (bottom panel). **h.** Single cell copy number heatmap (top panel) and subclone consensus copy number heatmap (bottom panel) from two merged wellDA-seq experiments.

### Technical performance of wellDA-seq

To assess the technical performance of wellDA-seq, we analyzed the ATAC and DNA data of MDA-MB-231 (MDA231) breast cancer cell line generated by two wellDA-seq replicate experiments (wellDA-1 and wellDA-2) and conducted a benchmarking analysis to other gold-standard unimodal technologies (**Supplementary Table 2**). We first compared the single cell ATAC (scATAC) data of wellDA-seq data to unimodal scATAC data generated from the same cell line (MDA231) using the 10X Genomics platform, and found that both wellDA-seq experiments show comparable transcription start site (TSS) enrichment scores and numbers of unique fragments between both datasets (**Fig. 1b**). We also observed a high correlation in the aggregate chromatin accessibility profiles between the wellDA-seq scATAC data and the 10X Genomics scATAC dataset (Pearson Correlation R = 0.91 and 0.92, p < 2.2e−16), as well as high correlations between the two wellDA-seq replicate experiments (Pearson Correlation R = 0.99, p < 2.2e−16) (**Fig. 1c-d).**

We next evaluated the scDNA-seq data quality of wellDA-seq relative to 4 other unimodal scDNA-seq methods (Arc-well^14^, 10X Genomics CNV^16^, DLP+^15^ and DOP-PCR^26^) by comparing two main quality control (QC) metrics: 1) ‘overdispersion’, the variations of read count distributions in each genomic bin, and 2) ‘breadth of coverage’, the uniformity of physical read coverage across the genome. To ensure a fair comparison, 120 randomly downsampled single cells from each method was used. We found that wellDA-seq datasets showed similar overdispersion metrics with Arc-well, but significantly lower overdispersion metrics compared to the other three scDNA-seq methods (p < 2.2e-16, Wilcox tests), suggesting the wellDA-seq datasets have very low levels of technical noise (**Fig. 1e**). Moreover, both wellDA-seq datasets showed significantly higher breadth of coverage compared to the 10X, DLP+ and DOP-PCR based unimodal methods (p < 2.2e-16, Wilcox tests) when using the same number of input reads per cell (750K randomly sampled), which indicating the superior read coverage uniformity across the genome in the wellDA-seq data (**Fig. 1e**). Finally, we estimated the number of single cells with wellDA-seq that passed quality control (QC) filters for both the ATAC and DNA modalities, which showed high percentages of overlapping cell (78.71% and 74.88%), indicating efficient performance in the amplification of both modalities in the experimental procedures (**Fig. 1f, Methods**).

To investigate the biological heterogeneity of MDA231, we merged the two wellDA-seq datasets and clustered the scATAC profiles and DNA copy number aberration (CNA) profiles independently. The co-clustering results from the two merged runs for both ATAC and DNA profiles were well integrated, indicating minimal bench effects (**Extended Data Fig. 2a**). Clustering of the DNA copy number profiles of 2,603 cells identified three superclones (S1, S2 and S3), which were further organized into 7 distinct subclones, whereas the clustering of the ATAC profiles revealed three main clusters (**Fig. 1g-h**). Mapping the three ATAC clusters directly to the DNA UMAP showed that cells from the ATAC clusters 1, 2 and 3 were from DNA superclones S2, S1 and S3, suggesting epigenetic heterogeneity in the ATAC profiles was reflected in the DNA copy number subpopulations at the superclone level (**Fig. 1g-h**). However, at the subclone level, the mapping of several subclones (C4-C7) within the ATAC cluster 2 were intermixed in the high-dimensional UMAP ATAC space, indicating that using the ATAC data only, we cannot resolve the subclone-level structure effectively (**Fig. 1g-h**).

By calculating the Pearson correlations between CNA profiles from scDNA data and ATAC-inferred CNA profiles of wellDA-seq, we found that the overall correlation was only a modest (median Pearson correlation R = 0.47) (**Methods, Extended Data Fig. 2b**). A more detailed comparison of the copy number heatmaps between the ATAC-inferred CNA profiles and the actual CNA profiles showed that while larger chromosomal events had some consistencies, there was also a very large number of false-positive CNA events reported in the ATAC-inferred CNA profiles (**Extended Data Fig. 2c**). Additionally, while the cell line data showed some agreement for larger chromosomal events, the data from the human breast tumor samples had very little concordance. Examples from three breast cancers (P3, P5 and P10) in which ATAC-inferred CNA could not accurately resolve any of the clonal substructure that was identified in the scDNA data and showed a larger number of false-positive CNA events, particularly in diploid regions (**Extended Data Fig. 2d**). These data show that computational methods to infer CNA profiles from scATAC data report inaccurate CNA information and substructure, which justifies the need to utilize co-assays such as wellDA-seq that can profile the genome and epigenomic experimentally as independent modalities.

Overall, these technical comparisons show that wellDA-seq can generate high-quality DNA and ATAC sequencing data simultaneously from the same single cells at high cellular throughput and high genomic resolution.

### Cell types with sporadic CNAs in normal breast tissues

Previous studies using genomic sequencing of bulk tissue samples have discovered that normal tissues in the human body, including breast tissues, harbor somatic genetic alterations^27–30^. However, these studies could not link these genetic events to specific cell type identities. Here, we applied wellDA-seq to normal breast tissues from two healthy female donors (P1, P2) and determined the specific cell types harboring somatic CNAs and their frequencies (**Supplementary Table 1 and 3**).

The first breast tissue (P1) was from a healthy young woman (late 20’s). The histopathological analysis of the Hematoxylin & Eosin (H&E) tissue sections confirmed normal cellular morphology and tissue structure (**Fig. 2a**). We profiled 1,339 single cells using wellDA-seq and identified 8 major clusters from the ATAC data (**Fig. 2b**). Based on the TSS-flanking ATAC fragments and the inferred RNA expression of the canonical cell type-specific marker genes^9,20^, the ATAC clusters corresponded to three epithelial cell types including LumSec, LumHR and Basal, two immune cell types (T-cells and myeloid cells), and three stromal cell types including fibroblasts (Fibro), endothelial cells (Endo) and pericytes (Peri) (**Fig. 2b, Extended Data Fig. 3a,b**). Unsupervised clustering on the single cell DNA data identified a rare cluster of 10 aneuploid cells in the tissue (0.75%), which all mapped to the LumSec clusters according their ATAC profiles (**Methods, Fig. 2b, c**). These aneuploid cells harbored a genomic gain of chr1q and different chromosome loss events at chr9, chr10q, and chr16q (**Fig. 2c, d**).

**Fig. 2.**
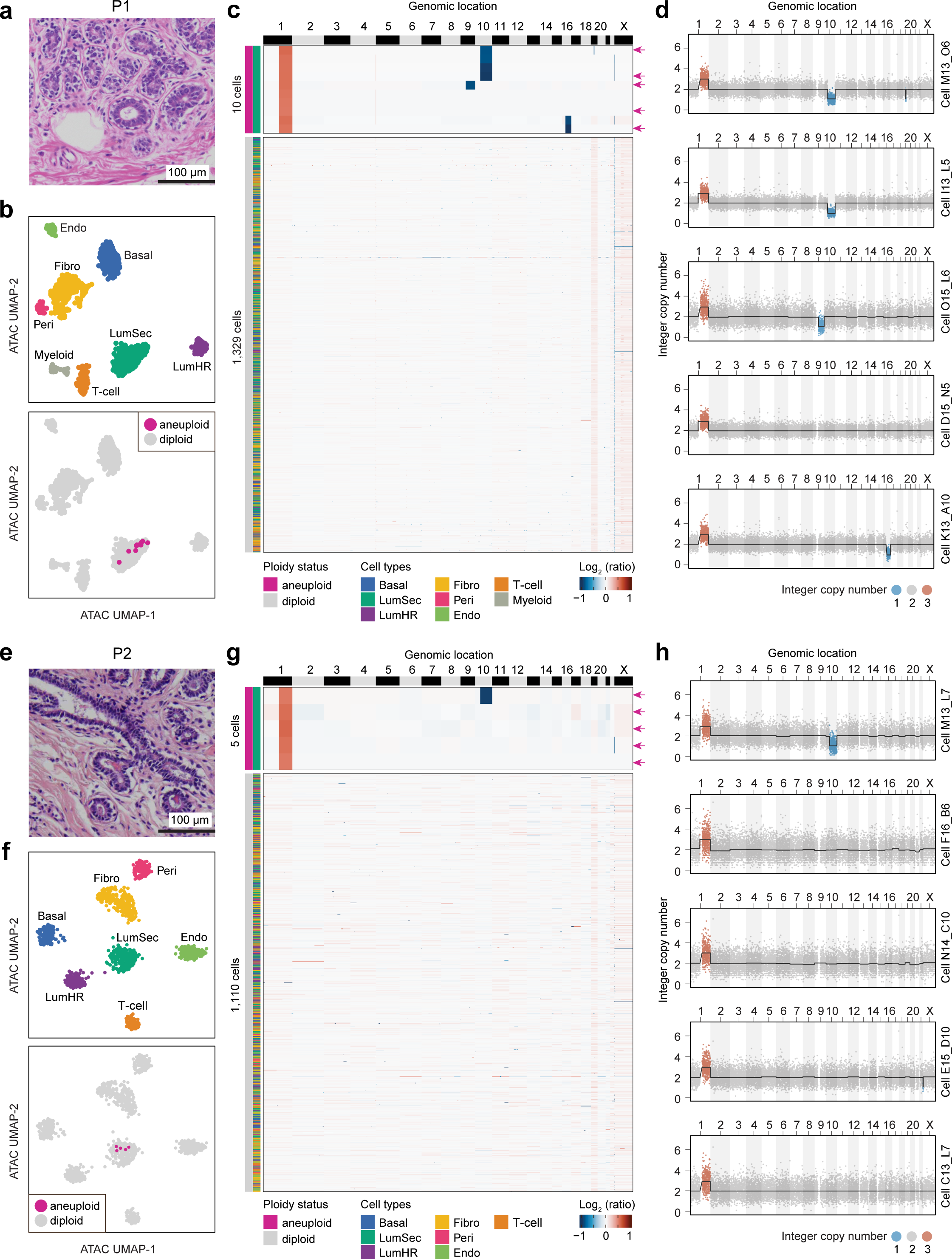
Cell types with sporadic copy number events in two normal breast tissues. **a.** Hematoxylin and eosin (H&E) image of the normal breast tissue histopathology from P1. **b.** UMAP of ATAC modality data of wellDA-seq showing clusters from 1,339 single cells from P1. **c.** Clustered heatmap of P1 showing the single cell CNA profiles of 10 aneuploid and 1,329 diploid cells. **d**. Plots of P1 showing the DNA data of wellDA-seq in 5 representative aneuploid cells indicated by the red arrows in panel **c** in the same order from top to bottom. **e.** H&E image of the normal breast tissue histopathology from P2. **f.** UMAP of P2 showing clusters of the ATAC modality data of wellDA-seq of 1,115 single cells. **g.** Clustered heatmap of P2 showing the single-cell CNA profiles of 5 aneuploid and 1,110 diploid cells. **h**. Plots of P2 showing the single cell DNA data of wellDA-seq in all the 5 aneuploid cells as indicated by the red arrows in panel **g** in the same order from top to bottom. In **a**, **e**, scale bar: 100 µm.

The second normal breast tissue (P2) was from a healthy older woman (early 40’s) (**Fig. 2e**). Analyzing the wellDA-seq data of 1,115 single cells identified 7 major cell types (**Fig. 2f, Extended Data Fig. 3a,b**). Consistent with P1, the CNA data in P2 also identified a rare subpopulation (0.45%) with 5 aneuploid cells (**Methods**). All of the 5 aneuploid cells had LumSec identities (**Fig. 2f, g**). The aneuploid cells all shared a chr1q gain and one additionally harbored a chr10q loss (**Fig. 2g, h**). Collectively, our wellDA-seq data from two disease-free women identified a rare population (<1%) of aneuploid cells harboring somatic CNAs in normal breast tissues. These aneuploid cells all had LumSec cellular identity and shared the same somatic chr1q gains and chr10q losses across the two patients.

***

### Copy number profiles of cell types and subclones in 9 breast cancers

We next applied wellDA-seq to profile the ATAC and DNA data of 19,669 single cells from 9 ER+ breast cancer samples including 3 cases of Ductal Carcinoma In Situ (DCIS) and 6 cases of invasive breast cancers (**Supplementary Table 1 and 4**). Unbiased high-dimensional clustering of the scATAC profiles from all 9 samples revealed 8 different cell types including LumSec, Basal, fibroblasts, endothelial cell, pericyte, myeloid, T cells and 9 LumHR-like clusters based on the top cell marker genes of cell types identified in the normal breast tissue samples (P1, P2) (**Fig. 3a, Extended Data Fig. 4a-b**). Analyzing the scDNA data identified 10,054 (51.11% of the total) single cells with copy number events, that were assigned as aneuploid cells. By mapping the ploidy status information to the ATAC UMAP, we found that most of the aneuploid cells were in the LumHR-like clusters, while a small number of aneuploid cells were from the T cell and pericyte clusters (**Fig. 3a**). By mapping the sample identifiers, we found that the aneuploid cells from different samples formed their own sample-specific clusters, whereas the diploid cells from different samples were well intermixed within each cluster (**Fig. 3a**). Across the 9 tumor samples, a mean of 2,185 (SEM = 197) cells were profiled and a mean of 1,088 (SEM = 298) cancer cells (i.e., aneuploid cells within the LumHR-like clusters) were identified. In majority of samples (8/9), the 8 normal cell types and 1 cancer cell population were detected (**Fig. 3b**).

**Fig. 3.**
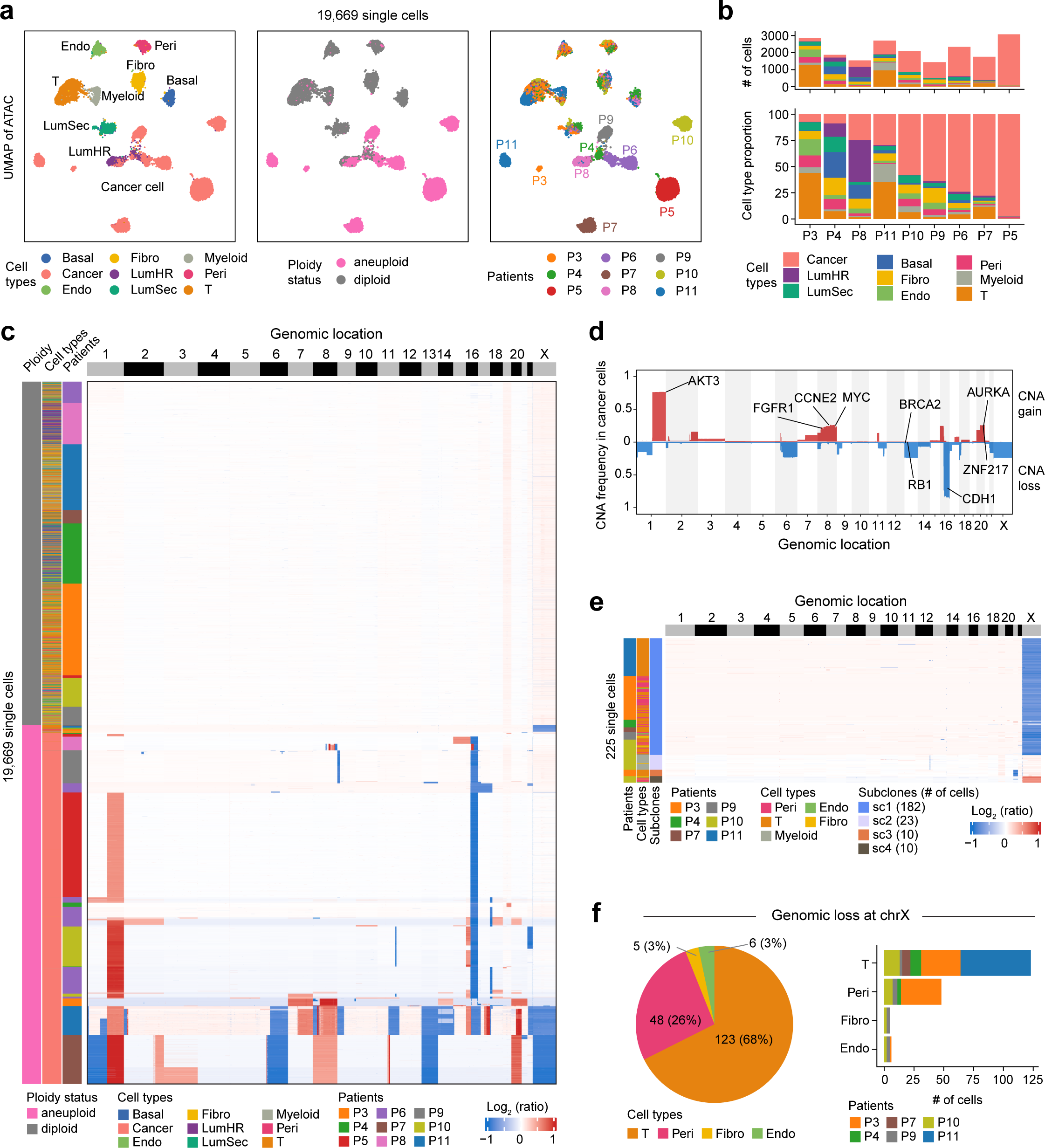
Overview of scATAC and scDNA data of 9 breast cancers profiled by wellDA-seq. **a.** Integrated UMAP of scATAC profiles of 19,669 single cells from 9 breast cancers, colored by cell types, ploidy status and patient. **b.** Cell number and cell type proportions profiled by wellDA-seq of each patient. **c.** Single cell DNA copy number heatmap of 19,669 single cells with left annotation bars indicating ploidy status, cell type, and patient. **d.** Copy number frequency of aneuploid cancer cells from 9 samples. **e.** Single cell DNA copy number heatmap of 225 aneuploid cells from non-epithelial cell types. **f.** Cell type proportion (left panel) and cell number distribution (right panel) of non-epithelial aneuploid cells with chrX loss.

We next integrated the single cell DNA copy number profiles of all 19,669 cells together with their cell type information and found that the normal diploid cells have ATAC profiles from the three normal epithelial cell types, three stromal cell types and two immune cell populations (**Fig. 3c**). In contrast, almost all of the aneuploid cells had ATAC profiles from the LumHR-like cells, indicating that they are cancer cells. By analyzing the copy number frequency across all of the cancer cells, we found that chr1q gain and chr16q deletions were the most frequent events in our data set, while chr8q gain, chr16p gain and chr20q gain were also observed in multiple patients (**Fig. 3d**).

In addition to the epithelial cells, we also detected aneuploid cells in a small number of the stromal and immune cells. Analysis of 225 aneuploid single cells from the non-cancer cells showed that most of these cells (80.88%) had a loss of chrX, while a small number of cells showed a focal loss of chr12, chr21 loss or a gain of chrX (**Fig. 3e, Extended Data Fig. 4c**). The chrX genomic loss mainly occurred in the T cells (68%) and pericyte (25%) and this observation was found in multiple patients (**Fig. 3f**). Among all the T cells that were sequenced (2,838), the X chromosome loss showed a frequency of 4.30%, but this event was not detected among the 911 myeloid cells that were profiled. This loss of chrX in T-cells identified by wellDA-seq is consistent with a previous study that reported chrX loss in T cells from colorectal cancer^31^. In addition to T-cells, our data also showed that chrX loss occurred in pericytes across multiple patients, at a frequency of 5.50% (49 out of 891 cells), which has not been previously reported. Collectively, these data show that aneuploid copy number rearrangements are mainly associated with epithelial cells, but that there are a few additional cell types in the tumor microenvironment (T-cells, pericytes) in which the X chromosome is lost and the resulting cells have expanded.

### Breast cancers ancestral cells and their epithelial cell-of-origin

To investigate the genetic alterations that associated with early clonal expansion and the epithelial cell-of-origin of breast cancer cells, we analyzed the 9 breast cancer samples profiled by wellDA-seq. In 6 of these samples (P4-P6, P8-P10), we directly detected rare ancestral cancer cell subpopulations using the scDNA profiles and revealed their epithelial cell type identity using the scATAC data.

In sample P10 (ER+, PR+, HER2-), clustering of the scDNA profiles of 1,203 single cells identified 6 different subclones. All subclones shared clonal CNA events including the chr1q gain, chr11q focal loss and chr16q loss. While a subset of the subclones also acquired additional gains on chr7 and chr16p, as well as chr 22 loss (**Fig. 4a**). Phylogenetic analysis of the consensus integer CNA profiles of each subclone showed that the initial normal (diploid) cell acquired 4 copy number events and formed subclone C6, which is the most recent common ancestor, then subsequently diverged and formed all of the other subclones (**Fig. 4a**). To determine the epithelial cell type identity of the ancestral subclone C6 and the other divergent subclones, we calculated the epithelial module score of their scATAC profiles (**Methods**), which showed that the ancestral C6 displayed the highest LumHR epithelial module score compared with LumSec and Basal, indicating that C6 originated from normal LumHR cells. Further analysis of the epithelial cell type identities of the other tumor subclones revealed that all of the subclones showed the highest score for LumHR, relative to their scores for the Basal and LumSec lineages. Notably, while the LumHR score was highest in all subclones, we also noticed a decreased in this epithelial identity in the more divergent subclones (C1-C5), suggesting that the cancer cells were losing their epithelial identity as they continued to evolve (**Fig. 4a)**.

**Fig. 4.**
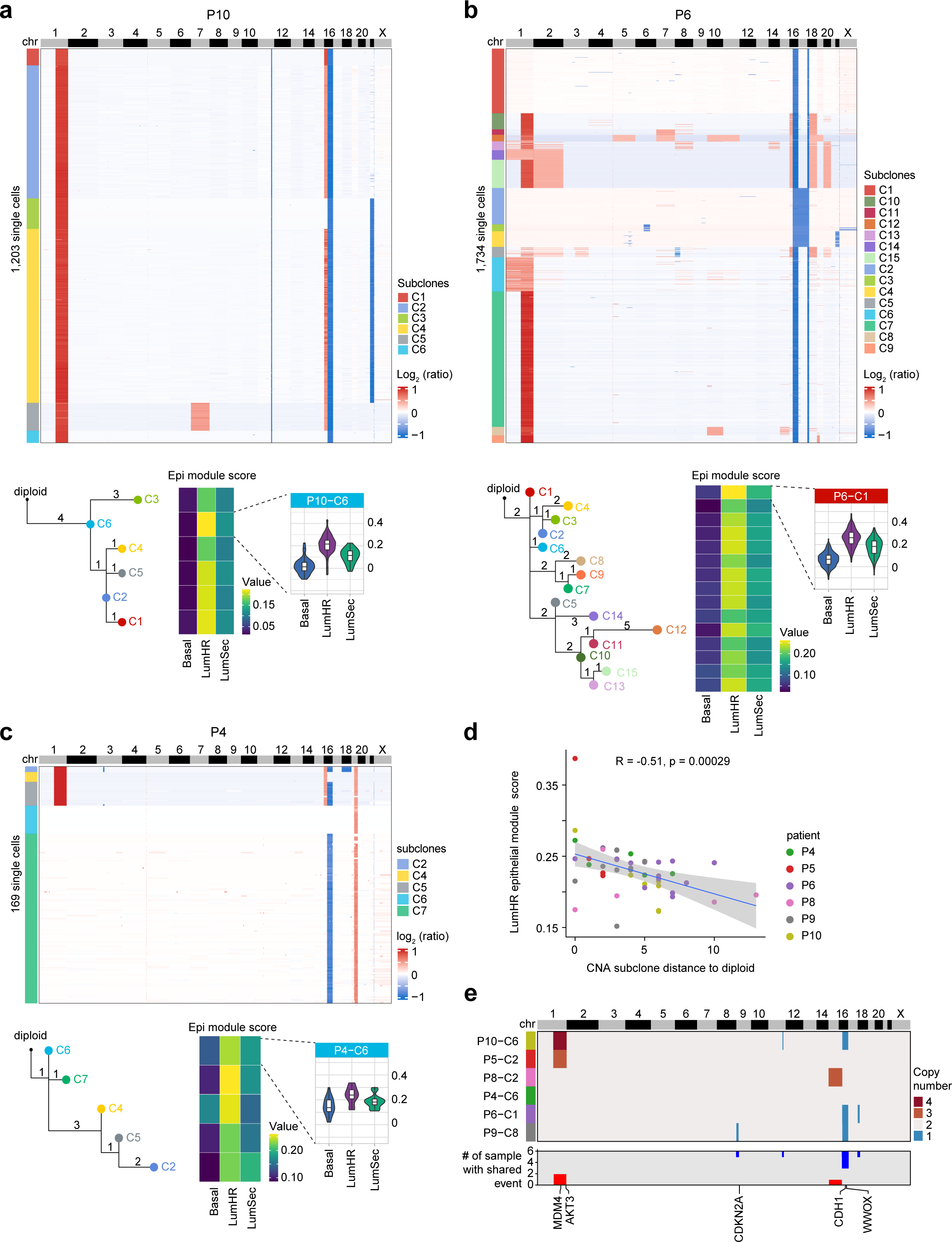
Genetic lineages and epithelial cell type identities of ancestral cancer cells. **a.** Heatmap of single cell copy number profiles of cancer cells from P10 with bottom left panel showing the phylogenetic tree constructed using subclonal consensus integer copy number profile, and right bottom panel showing the epithelial module scores for each cell type in each subclone. **b.** Heatmap, phylogenetic tree and subclone epithelial cell type module scores for P6. **c.** Heatmap, phylogenetic tree and subclone epithelial cell type module score for P4. **d.** Pearson correlation coefficient between phylogenetic distances to diploid for each aneuploid subclone and the LumHR epithelial cell type module score of each subclone in 6 breast cancer samples. **e.** Consensus integer copy number profiles of the ancestral cell subclones from 6 breast cancers, with bottom panel showing the number of samples with shared CNA events.

In another ER+ and PR+ sample (P6), clustering of the scDNA data of 1,734 single cells identified 15 subclones. Phylogenetic analysis identified C1 as the ancestral subclone that harbored chromosome losses of chr16q and chr18p. Similar to P10, the scATAC data indicated that the ancestral populations originated from the LumHR epithelial cells, and continued to lose their of epithelial cell type identity during cancer progression in the more divergent subclones (C2-C15) (**Fig. 4b**). In samples P5 and P8, we identified similar patterns, in which the ancestral cancer cell populations were also from the LumHR epithelial cells, and lost their epithelial identities during cancer evolution (**Extended Data Fig. 5a-b**). Similarly, in P4 and P9, the ancestral cancer cells were also from the LumHR epithelial cells. While the ancestral cells did not have the highest LumHR scores compared to the other tumor subclones in these two patients, there was a consistent trend of losing the LumHR cell type identity as the subclones acquired additional CNAs during cancer evolution (**Fig. 4c, Extended Data Fig. 5c**).

We further investigated the relationship between the distance of subclones in the clonal lineage from the diploid cells and the LumHR epithelial score in all 6 samples. This analysis revealed a negative relationship (R = -0.51, p = 0.29 × 10^-3^) between the CNA distance and the LumHR epithelial score, which showed that the LumHR epithelial score was decreased as the cancer subclones evolved and progressed in our dataset (**Fig. 4d**). In addition, by summarizing of the CNA events occurring in all of the ancestral cells across the 6 samples, we found that chr16q loss occurred in 3 samples, while chr1q gain occurred in 2 samples, suggesting they are early events during ER+ breast cancer progression (**Fig. 4e**). Taken together, these data identified ancestral cancer cell populations in ER+ breast cancers and showed that they originated from the LumHR epithelial cells, with decreasing lineage identity and dedifferentiation during cancer progression.

### Impact of subclonal copy number alterations on chromatin accessibility

We analyzed the wellDA-seq data of the cancer cells from 9 breast tumors to quantitatively investigate the effect of subclonal CNAs on chromatin accessibility. In patient P8, wellDA-seq identified 4 different subclones C1-C4 based on the CNAs profiles (**Fig. 5a**). Phylogenetic analysis of consensus subclone CNA profiles showed that C1 and C3 resided in a common branch of the CNA tree, while C2 and C4 were grouped in another branch (**Fig. 5b, Methods**). Consistent with this result, clustering on the scATAC data showed that C1 and C3 co-clustered, while C2 and C4 were clustered together, indicating a high concordance between genomic CNAs and chromatin accessibility (**Fig. 5c**). To estimate the impact of subclonal CNAs on chromatin accessibility, we compared C1 and C4 as they harbored distinct genotypes and were abundant in the tumor. Between C1 and C4, we identified 780 subclonal genomic bins (SGBs), which included many focal events on chr8 and chromosome arm-level events on chr15 and chr16 (**Fig. 5d-e, Methods**), and identified 36 differentially accessible peaks (DAPs) that located in 29 genomic bins, of which 28 (96.55%) genomic bins were SGBs (**Fig. 5d, Methods**).

**Fig. 5.**
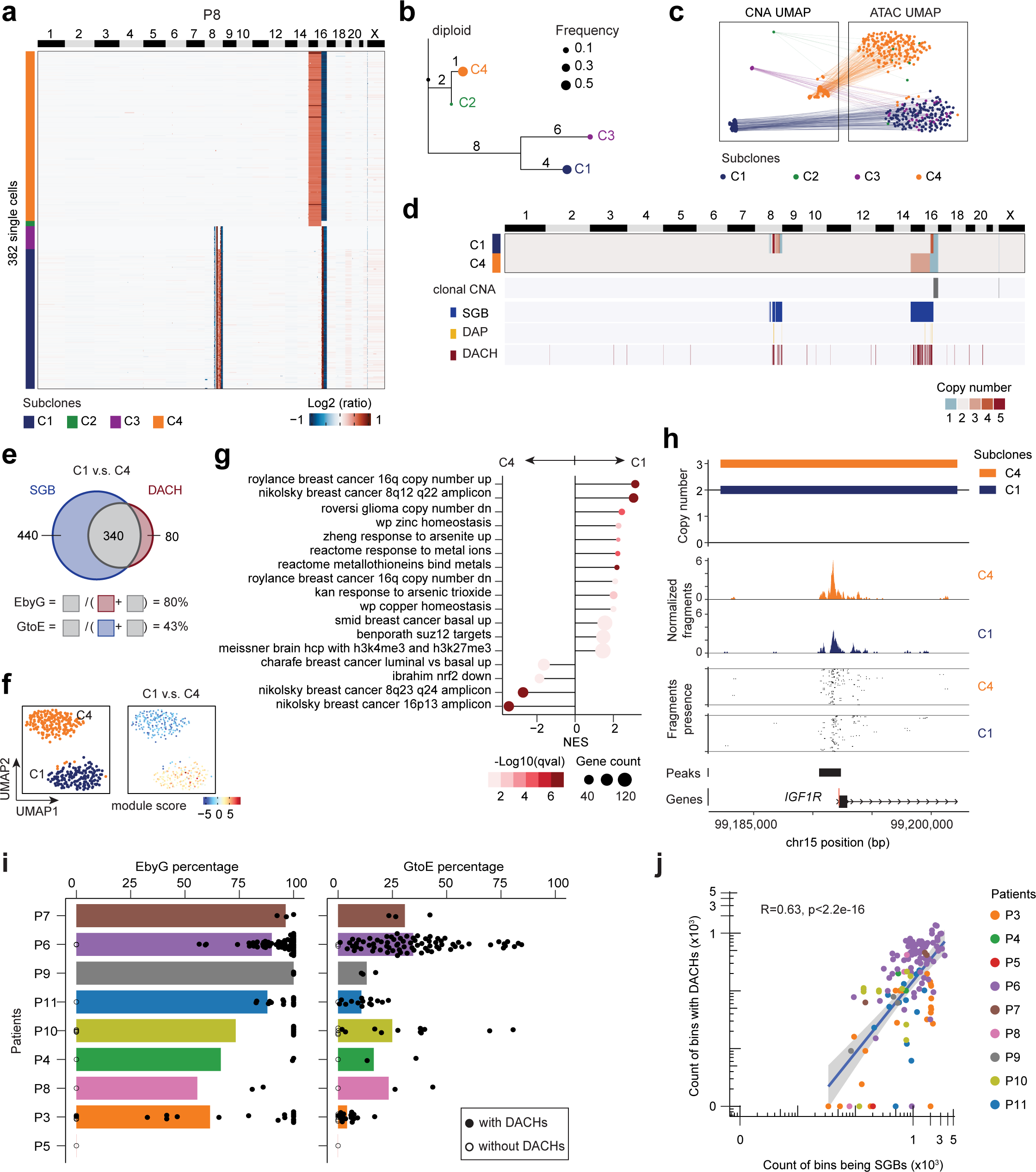
Impact of subclonal copy number alterations on chromatin accessibility. **a.** Clustered heatmap of P8 showing the CNA profiles of 382 single cells grouped by subclones. **b.** MEDICC2 phylogenetic tree of consensus subclones, with node size denoting the cellular frequency of each subclone. **c.** 382 single cells being projected from CNA UMAP space to ATAC UMAP space. **d.** Heatmap showing the consensus integer copy number profiles of subclone C1 and C4, with bars below for clonal CNA events, SGB, DAP, and DACH annotations. **e.** Venn diagram showing the overlap of the SGB and DACH events in a comparison of C1 and C4. **f.** UMAP of ATAC data showing the single cells colored by the subclones C1 and C4 (left panel) and the module scores of C1-specific DACH events (right panel). **g.** Lollipop plot showing the significant gene signatures that were enriched for the DACH events between C1 and C4. **h.** Track plot showing the integer copy number, normalized ATAC fragments, presence of Tn5 insertions in 50 randomly selected cells, and the peaks in the 200 kilobase window around the transcriptional start site of the gene *IGF1R*. **i.** Barplot showing the average EbyG and GtoE percentages in each patient sample. **j.** Scatter plot comparing the count of bins with DACH and SGB with the Pearson correlation coefficient and p-value labelled. In **i** and **j**, each dot denotes a subclone-to-subclone comparison.

To overcome the sparsity of scATAC, we performed the *cis-*Co-accessibility Networks analysis using Cicero^32^ to identify chromatin hubs in single cells and further determined the differentially accessible chromatin hubs (DACHs) between C1 and C4 (**Methods**). We found that 420 genomic bins containing DACHs (**Fig. 5d-e**, **Methods**), of which 340 DACH-overlapped with SGBs (**Fig. 5d-e**). We then computed how this number was relative to the total SGBs number, which we refer to as ‘GtoE’ percentage, and also how it was relative to the number of total DACH-overlapped genomic bins, referred as ‘EbyG’ percentage (**Fig. 5e, Methods**). In the comparison of C1 and C4, the GtoE percentage was intermediate showing that 43% genomic CNA events (340 of 780 SGBs) affected the epigenetic chromatin accessibility *in cis*. In contrast, the EbyG percentage was high showing that 80% of genomic bins with DACHs (340 of 420 bins) resided in SGBs, suggesting that the majority of the chromatin accessibility differences between C1 and C4 were a result of *cis-*chromatin changes in subclonal CNAs (**Fig. 5e-f**). We next leveraged the MSigDB gene sets^33^ and asked which gene sets were associated with the DACHs. By performing the module-based gene enrichment analysis as previously described^20^, we found that these DACHs were enriched for many gene sets related to gene perturbations in chr8 and chr16 in breast cancer, which were consistent to many subclone-specific SGBs (**Fig. 5d,g; Methods**). We also examined the known breast cancer genes^34^ residing in the DACH-overlapped SGBs. For example, we found that *IGF1R* showed fewer copy numbers in C1 compared to C4, while the normalized ATAC fragment counts were concordantly lower in C1 than C4 (**Fig. 5h**).

Next, we analyzed 142 subclone-to-subclone comparisons in all patient samples (**Methods**). We found that most comparisons (128/142) had DACHs between subclones, while some comparisons did not. The absence of DACHs between two subclones were not (p > 0.05, Wilcoxon) associated with technical factors (e.g., lower cell number or ATAC fragment number), but were associated with smaller phylogenetic distances (p < 0.05, Wilcoxon) (**Extended Data Fig. 6e-g**), indicating a correct GtoE estimated by wellDA-seq. Taken together, we found that the EbyG percentages were on average 82.27% (SEM = 2.59%), suggesting that a large percentage of chromatin changes were directly linked to CNAs in *cis*. While we found that the GtoE percentages exhibited a large range across patients and were on average 25.12% (SEM = 1.85%), indicating that not all genomic CNAs necessarily elicited *cis*-associated changes of chromatin accessibility (**Fig. 5i, Extended Data Fig. 6a-d**). Last, there was a positive correlation between the count of SGBs and DACHs (R = 0.63, p < 0.05), suggesting that an increased genomic instability is associated with pervasive chromatin remodeling (**Fig. 5j, Methods**).

### Genetic hardwiring and epigenetic plasticity in clonal lineages

To investigate whether cancer phenotypes are genetically hardwired, or alternatively plastic across subclones, we studied the relationship between the copy number lineages and the chromatin accessibility programs of cancer subclones across the 9 breast tumors.

In the patient P6, we identified 15 subclones that shared a common lineage with clonal events such as 16q loss and 18p loss (**Fig. 6a**). Additionally, these subclones had many divergent subclonal events (e.g., 1q gain and 18 gain) indicating a high subclonal diversity in this sample. By projecting the DNA subclone to the scATAC clustering results, we found that not all of these subclones were mapped to well separated ATAC clusters. For example, the subclones C13, C14 and C15 were co-mapped to the same ATAC cluster despite the large chromosomal arm-level CNA events on chr1 and chr2 (**Fig. 6a,b)**. Using the Pearson correlation coefficient between the CNA-based and ATAC-based pair-wise distances of subclones, we found that the global concordance between the subclonal CNAs and chromatin accessibility profiles in P6 was intermediate (0.48) (**Fig. 6c, Methods)**. To identify specific cancer phenotypes (epigenomic programs) that were heritable or plastic in the tumor, we leveraged the gene signatures from the MSigDB cancer hallmark database^15^ and assessed the extent to which these phenotypes in the tumor followed the structure of the phylogenetic tree (**Methods**). For each gene signature, we calculated the ATAC-based distances of the subclones and computed how these distances were correlated with the genomic CNA-based Manhattan distances to quantify the level of heritability. A high correlation score suggested a heritable phenotype showing comparable expression levels in the neighboring subclones in the phylogenetic tree, whereas a low correlation score implied a plastic phenotype showing expression levels independent of genetic lineages (**Methods**). In this sample, we found that the top genetically heritable programs included TNF-α signaling via NF-kB, inflammatory response, and apical junction, whereas the top plastic gene sets included TGF-β signaling, angiogenesis, and peroxisome. (**Fig. 6d**). In another sample P10, we identified 6 subclones that shared clonal events of 1q gain and 16q loss, in addition to having divergent CNA subclonal events at chromosome 7 and 16p (**Fig. 6e**). Overall, the global CNA-ATAC concordance in P10 was also intermediate (0.55), reflecting that the subclones did not map to distinct ATAC clusters (**Fig. 6f, g**). We found that the genetically hardwired programs included apical junction, glycolysis, and reactive oxygen species pathway, while the plastic signatures included fatty acid metabolism, hypoxia, and interferon gamma response (**Fig. 6h**).

**Fig. 6.**
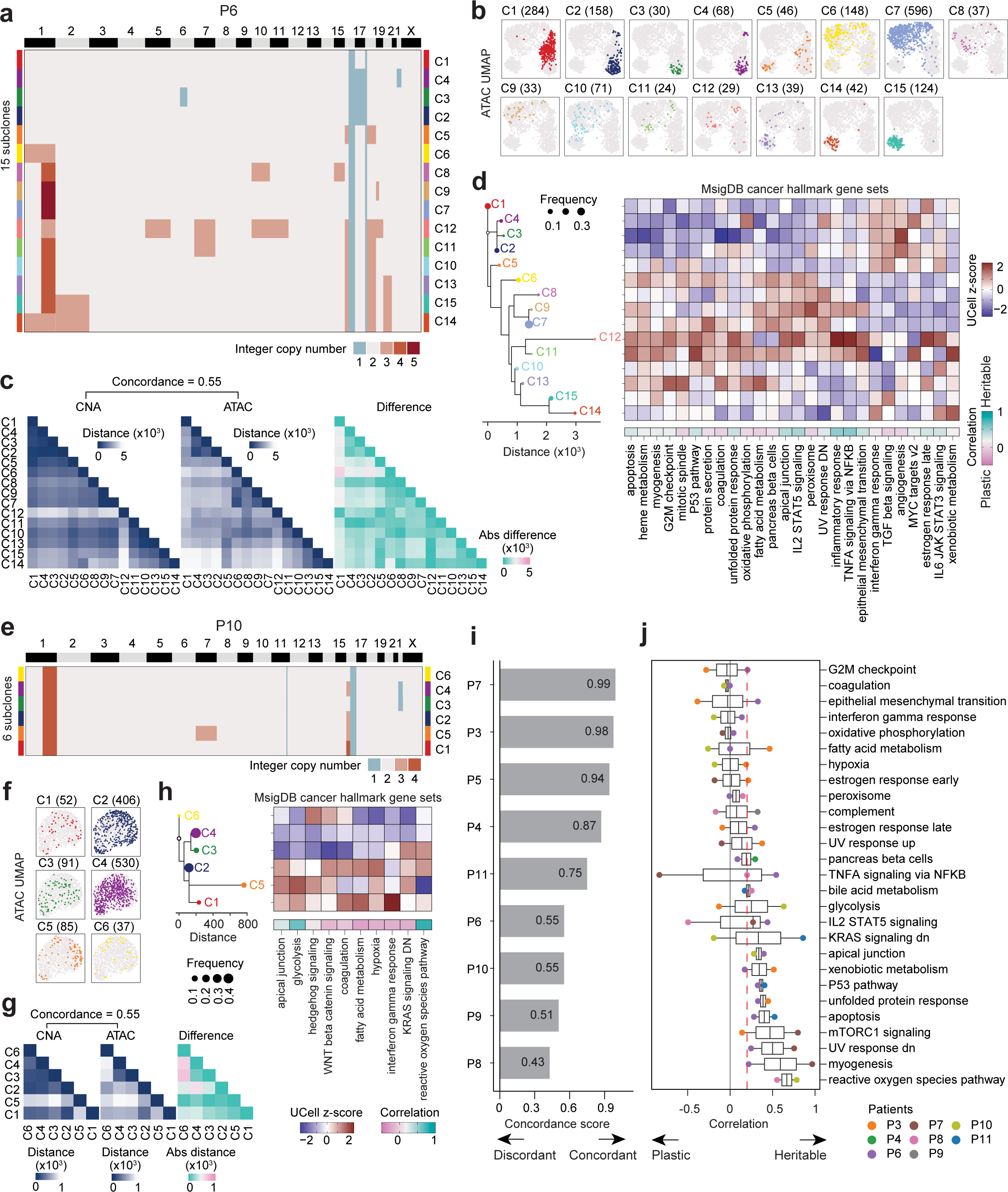
Genetic hardwiring and epigenetic plasticity in breast tumors. **a.** Heatmap of P6 showing the consensus integer copy number profile of the 15 subclones. **b.** UMAP plots of ATAC data from P6 colored by cells from DNA subclones. **c.** Heatmap plots of P6 showing the subclone-to-subclone distance matrix based on the CNA and ATAC data, along with the absolute difference of these two matrices used to calculate the global concordance score. **d.** Minimum evolution tree of clonal lineages (left panel) and heatmap (right panel) in P6 showing the relative gene activities of the cancer hallmark gene sets. The color bar beneath the heatmap represented the correlation metric quantifying the level of heritability and plasticity for each gene signature. **e.** Heatmap of P10 showing the consensus integer copy number profile of the 6 subclones. **f.** UMAP plots of ATAC data from P10 colored by cells from DNA subclones. **g.** Heatmap plots of P10 showing the subclone-to-subclone distance matrix based on the CNA and ATAC data, along with the absolute difference of these two matrices. **h.** Minimum evolution tree of clonal lineages and heatmap in P10 showing the relative gene activities of the cancer hallmark gene sets with heritability and plasticity scores below. **i.** Barplot showing the concordance scores between CNA profiles and chromatin accessibility profiles in each patient sample. **j.** Boxplot showing the heritability/plasticity scores for recurrent signatures colored by patient.

This analysis was also performed on the 7 other samples showing that the concordance score was on average 0.73 (SEM=0.07) with a range of 0.43 to 0.99 across samples (**Fig. 6i; Extended Data Fig. 7**). The concordance scores were not associated with technical factors such as cells count, subclones number, and ATAC fragment number (**Extended Data Fig. 8**). Moreover, our wellDA-seq data showed that in these 9 tumors, the reactive oxygen species, myogenesis, apoptosis and P53 pathways were among the top heritable gene signatures, while the G2M checkpoint, coagulation, epithelial-mesenchymal-transition (EMT), and interferon gamma response were top plastic gene signatures, suggesting that cancer cells from different genetic lineages adopted similar phenotypes of modulating their surrounding tumor microenvironment (**Fig. 6j**).

## Discussion

In this study, we report wellDA-seq, the first high-genomic resolution and high-throughput single cell whole genome and chromatin accessibility co-assay method. By applying this method to study genotype-phenotype interactions in cancer cells from 9 ER+ breast cancers, we made important biological discoveries regarding the epithelial cell-of-origin of ER+ breast cancer, quantifying the impact of genetic variations on epigenome changes, and identified genetic hardwiring and plasticity in clonal lineages that would be challenging to resolve with unimodal methods.

A major challenge in breast cancer evolution field is to identify ancestral subpopulations and their epithelial cell-of-origin. Here, using wellDA-seq we directly detected the ancestral cancer cell populations in 6 ER+ breast cancers, which showed that they all originated from the LumHR epithelial cell types. In 6 samples with ancestral cancer cells, we found that chr16q loss occurred in 3 patients and chr1q gain was detected in 2 patients, implying that these may be early CNA events involved in tumor initiation and early expansion in these patients. The wellDA-seq results enabled us to directly map the epithelial cell type chromatin profiles to the phylogenetic trees, which revealed that the LumHR signature decreased during cancer evolution, suggesting a progressive loss of the original epithelial cell type identities and is consistent with dedifferentiation in cancer.

Our study investigated another fundamental question: do subclonal CNA events have a direct impact on chromatin remodeling and how much chromatin accessibility changes were caused by subclonal CNA events. Using wellDA-seq, we quantify that about 82.27% of chromatin accessibility changes between subclones were impacted by the *cis-*effects of CNAs. Alternatively, our data also showed that a small number of chromatin accessibility changes (17.73%) were associated with *trans-*effects of CNAs. The *trans-*chromatin changes may be due to *cis-*chromatin changes at transcription factors or epigenetic regulators that resulted in broad changes of chromatin accessibility at different sites across the genome.

Another important question wellDA-seq addressed is that wellDA-seq provided directly evidence that the genetic hardwiring and phenotype plasticity presented between tumor subclones and modulated tumor progression. By applying wellDA-seq to 9 breast cancers, we reconstructed clonal lineages and investigated how cancer phenotypes mapped to the genetic lineages. Our global analysis found that about half of the breast tumors were highly concordant in their genetic lineages and epigenomic profiles, while the other cases showed many differences in the chromatin profiles and their relationships to the genetic lineages. By investigating specific cancer phenotypes across patients, we found that the top cancer phenotypes that were genetically heritable included reactive oxygen species, apoptosis and P53 pathway, while the top plastic gene signatures included G2M checkpoints, EMT, and interferon gamma response. Indeed, previous data has shown that EMT is often a plastic phenotype with many transition states in cancer cells and has not been linked to specific genetic mutations that expand in a tumor^35,36^.

The capability of simultaneously capturing both genotype and phenotype of single cells also makes wellDA-seq identify non-epithelial cell types (e.g., T-cells and pericytes) with somatic CNA events, such as chrX deletions in breast cancers, suggesting that somatic genomic aberrations were also presented in the tumor microenvironment. In addition to cancer samples, by applying wellDA-seq to normal breast tissues of two healthy women, we identified rare aneuploid cells (with frequency <1%) harbored somatic CNAs (e.g., chr1q gain and chr10q loss) and showed the LumSec cell identity. These applications of wellDA-seq lay the groundwork for larger future studies that could include many tissue types to understand their prevalence across different organ systems, such as the NIH Somatic Mosaicism across Human Tissues (SMaHT) network.

Our study has a few limitations. First, our samples were limited to a relatively small cohort size that included 2 normal breast tissues and 9 ER+ breast cancer patients. Therefore, our study serves as a proof-of-concept, and the biological discoveries we have reported may (or may not) be generalizable to larger cohorts of patients. Second, we applied wellDA-seq primarily for DNA copy number and chromatin accessibility analysis, which involved sequencing the whole genomes of single cells at sparse coverage depth and is not amenable for mutation detection. An important future direction will be to perform target capture of the DNA libraries from wellDA-seq and to achieve higher coverage depth to detect mutations at base-pair resolution.

In closing, wellDA-seq provides a highly scalable approach to perform single cell whole genome and chromatin accessibility sequencing simultaneously from single cells. In future studies, the application of this method holds great potential for unraveling the complex interactions between genetic events and epigenomic programs of single cells. We expect that wellDA-seq will have broad applications in fields as diverse as immunology, developmental biology, aging research, neuroscience, and cancer research, to improve our understanding of human diseases and normal tissue biology.

## Data and materials availability

The DNA and ATAC data from this study has been deposited to the Sequence Read Archive (SRA): PRJNA1081815 and Gene Expression Omnibus (GEO): GSE260864, respectively. The code can be found at https://github.com/navinlabcode/wellDA-seq.

## Acknowledgements

This work was supported by grants to N.E.N. from the NIH National Cancer Institute (RO1CA240526, RO1CA236864). N.E.N. is an AAAS Fellow, AAAS Wachtel Scholar, Damon-Runyon Rachleff Innovator, Andrew Sabin Fellow, and Jack & Beverly Randall Innovator. Y.Y. was supported by the GSBS Investing in Student Futures Fellowship, Larry Deaven Fellowship, and Ray Meyn Scholarship. We thank Yuehui Zhao, Nicolass Baudoin, Jerome Lin, Anna Casasent, Stacey Carter, Betsy Bonefas, Ivan Marin, and Margarita F Rojas-Barrett for their help.

## Author Contributions

K.W. designed and developed the wellDA-seq method with help from H.E. and Y.Y.; K.W., H.E., J.L. performed the experiments with help from S.B., Z.X., Y.L. and E.S.; Y.Y. and K.W. performed the analysis with help from C.T., and J.W.; J.M., C.N. and A.T. provided clinical samples to support these studies. N.N. and K.W. supervised the project. K.W., Y.Y. and N.N. wrote the manuscript.

## Methods

### Cell lines

The human breast cancer cell line (MDA-MB-231) were obtained from the MD Anderson Characterized Cell Line Core Facility and were cultured in Dulbecco’s Modified Eagle’s Medium-high glucose (DMEM, Sigma, D5976) supplemented with 10% Fetal Bovine Serum (FBS) (Sigma, F0926), 2mM L-Glutamine and 1% Penicillin/Streptomycin (Sigma P4333). Cells were passaged every 2–3 days and seeded at approximately 500,000 cells/mL. The cells were maintained at 37°C in 5% CO2 incubator before running the experiments.

### Human breast tissue samples

Breast tissues from healthy women were collected from reduction mammoplasties, while breast cancer tissues were collected as fresh tissues after the breast cancer surgery. To dissociate the single cells^9^, breast tissues were minced and incubated in 1 X dissociation buffer (1.875% BSA fraction V (Gibco, 15260037) and 1g/ul Collagenase A (Sigma, 11088793001) in DMEM F12/HEPES (Gibco,113300)) for 3 hours with rotating in 37°C incubator, then treated with trypsin containing 0.25% EDTA (Corning, 25053CI) for 5 min and neutralized with DMEM (Sigma, D5796) containing 10% FBS followed by filtration through 70 μm filters. After centrifugation, cell pellets were incubated with 1 × MACS RBC lysis buffer (MACS,130-094-183) for 10 min at room temperature to remove red blood cells. Cell pellets were then collected and washed by DMEM. Single cell solutions were preserved as viable cell suspensions by cryofreezing 10% DMSO and 90% FBS freezing media until thawed for the experiments. This study was approved by the Institutional Review Board (IRB) under Baylor College of Medicine (H-46622 and H-44538).

### Tn5 transposome assembly

wellDA-seq utilizes two different Tn5 transposome loaded with two different sets of adapters. The first Tn5 (Tn5-1) used to label open chromatin regions was assembled as below: custom adaptors preparation: 100 µM oligonucleotides with Tn5 mosaic end (ME) sequences (DEpi_ME_t1: 5’-GCCTCCCTCGCGCCATAGATGTGTATAAGAGACAG-3’ and DEpi_ME_t2: 5’-CTTGCCAGCCCGCTCAGAGATGTGTATAAGAGACAG-3’) were separately incubated with 100 µM ME reverse oligonucleotide (5’-/phos/CTG TCTCTTATACACATCT-3’) at 95°C for 3 mins then allowed to cool to 25° gradually (0.1°C per second) and incubated for an additional 5 mins. The annealed adaptors were then diluted to 25 µM and incubated with equal volumes of Tn5 enzyme (Diagenode, Tagmentase C01070010-20) and glycerol at 37°C for 10 mins followed by 30 mins at 25°C. The second Tn5 used to label DNA regions was purchased from Illumina (20034198).

### wellDA-seq procedure

#### Step 1: Cell preparation for bulk nuclei tagmentation

Cells were collected from plates (MDA-MB-231) or cryovials (human normal and tumor breast tissues), washed with 1X Dulbecco’s Phosphate Buffered Saline (DPBS, D8537, Sigma) containing 0.04% BSA and passed through a 40 μm Flowmi Cell Strainer. Around 500k cells were incubated with chilled lysis buffer (10 mM Tris-HCl (pH 7.4), 10 mM NaCl, 3 mM MgCl_2_, 0.1% Tween-20, 0.1% Nonidet P40 substitute, 0.01% Digitonin and 1% Bovine Serum Albumin (BSA)) for 4-5 mins to permeabilize cell membrane. Cells were then pelleted and washed using chilled Wash Buffer (10 mM Tris-HCl (pH 7.4), 10 mM NaCl, 3 mM MgCl_2_, 0.1% Tween-20, and 1% BSA). Cells were centrifuged at 500g for 5 min at 4°C, followed by careful removal of the supernatant. After resuspending the cell pellet in DPBS, fluorescent dye DAPI (4′,6-diamidino-2-phenylindole) was added to count nuclei using a Countess II FL Automated Cell Counter (Life technologies, AMQAX1000). Bulk tagmentation was performed in a 50 μl reaction containing: 25 μl 2 × TD buffer, 1 μl 0.5% digitonine, 5 μl assembled Tn5 (Tn5-1), 14 μl DPBS with nuclei (30k-35k) and 5 μl nuclease-free water. The reaction was incubated at 37°C for 30 min with intermittent shaking (850 rpm, 15s mix and 15s pause). To increase the cell number for later on-chip dispensing, 2-3 tagmentation reactions were performed. The cell suspension of transposed nuclei was centrifuged, and the pellet was resuspended with 0.5 × DPBS containing DAPI.

#### Step 2: On-chip whole genome transposition and cell barcoding

(1) Cell dispensing and nanowell selection

Option 1: One round of cell dispensing and imaging. Tagmented cell suspensions with final concentration of 1.05 cell/35 nl were dispensed into ICELL8 350v nanowell chips (35nl/nanowell) using ICell8 cx single cell dispensing system (Takara Bio). The chips were then imaged with DAPI configuration and only nanowells containing a single cell were selected by the CellSelect Software (Takara Bio) for downstream dispensing steps.

Option 2: Two rounds of cell dispensing and imaging. Tagmented cell suspensions with final concentration of 1.05 cell/25nl were dispensed into ICELL8 350v nanowell chips (25 nl/nanowell) with the following modified settings: change ‘Dispense VolumeNL’ from 35 to 25 under Volume 35nL and VolumeSample35 under menu of service/configure/Biodot configuration/volumes, then press “Done” under Utilities/Application manger. The chips were then imaged with DAPI configuration to identify empty wells and wells containing cells. Then, 25 nl of cell suspensions were dispensed into empty wells while 25 nl of 0.5 × DPBS were dispensed into wells containing cells. The chips were then imaged again with DAPI configuration and only nanowells containing one cell were selected for downstream dispensing steps.

(2) Cell lysis: 35 nl lysis buffer (200 μl recipe: 180 μl lysis buffer (30 mM Tris-HCl, ph8.0, 5% Tween, 0.5% TritonX-100), 20 μl protease (1.36 AU/ml)) was dispensed into the selected nanowells with single cells. The nanowell chip was then sealed with RC film (R5275, Takara Bio), centrifuged at 1000 g at 4°C for 5 min then incubated at 59.7°C 5 s, 54.5°C 30 min, 79°C 11 s, 75.3°C 15 min with lid 72°C.

(3) Second tagmentation: 35 nl tagmentation mixture (160 μl recipe: 148-152 μl 2 × TD buffer, 12-8 μl TDE1, Illumina) was dispensed into the selected nanowells of the chip. Then incubated the chip at 59.7°C for 5 s, 54.5 °C 10 min.

(4) Neutralization and first index dispensing. 35 nl of neutralization mixture with different PCR index (20 μl recipe for each index: 10 μl 5 × Kapa Hifi Fidelity buffer, 4 μl 10 mM dNTP, 1 μl 0.5 M EDTA, 4 μl H_2_O, 0.5 μl 100 μM wellDA_DNA_S5XX: 5’-AATGATACGGCGACCACCGAGATCTACACXXXXXXXXTCGTCGGCAGCGTC-3’, and 1.5 μl 100 μM wellDA_ATAC_S5XX: 5’-AATGATACGGCGACCACCGAGATCTACACTGAXXXXXXXXGCCTCCCTCGCGCCAT-3’, XXXXXXXX represent 8 bp index sequence) (**Supplementary Table 4-5**, 72 ATAC-CB1 and 72 DNA-CB1 primers) was dispensed into the selected wells. The chip was centrifuged and incubated at 54.9 °C for 5 s, 49.4 °C for 30 min, 4 °C hold.

(5) Second index dispensing. 35 nl of second indexing mixture (20 μl recipe for each index: 10 μl 5× Kapa Hifi Fidelity buffer, 0.348 μl 1 M MgCl2, 7.652 μl H_2_O, 0.5 μl 100 μM wellDA_DNA_IN_N7XX: 5’-CTGAGTCGGAGACACGCAXXXXXXXXGTCTCGTGGGCTCGG-3’, and 1.5 μl 100 μM wellDA_ATAC_N7XX: 5’-CAAGCAGAAGACGGCATACGAGATXXXXXXXXCTTGCCAGCCCGCTCAG-3’, XXXXXXXX represent 8 bp index sequence) (**Supplementary Table 4-5**, 72 ATAC-CB2 and 72 DNA-CB2 primers) was dispensed into the selected nanowells.

(6) PCR master mix. 35 nl PCR master mix (200 μl recipe: 40 μl 5 × Kapa Hifi Fidelity buffer, 40 μl KAPA HiFi HotStart DNA Polymerase (1 U/μl), 120 μl H_2_O) was dispensed into each selected nanowell. After centrifugation, the chip was incubated following the PCR cycles of 72.1°C 8 min, 99.6 °C 30 s, 12 × (99.6 °C 20 s, 57.5 °C 5 s, 62.7 °C 30 s, 72.1 °C 1 min), 72.1 °C 2 min, 4 °C hold.

(7) PCR product collection and purification. The PCR product was then collected using Collection Module (Takara Bio) by centrifuging the chip facing downwards at 3,220 g for 5 min at 4 °C and purified with 1.5 × Ampure XP beads (Beckman), followed by DNA QC and trace checking.

#### Step 3: Off-chip DNA and ATAC libraries enrichment

The purified library was then used to enrich ATAC and genomic DNA fragments independently using primers complementary to respective unique adapters. For DNA library amplification, 30 ng of purified product is mixed with 25 μl 2 × KAPA HiFi HotStart ReadyMix (KK2602, Roche), 1.5 μl forward primer (Bioo-PCR-F: 5’-AATGATACGGCGACCACCGAGATCTACAC-3’, 10 µM), 1.5 μl reverse primer (wellDA_DNA_OUT_N7XX: 5’-CAAGCAGAAGACGGCATACGAGATXXXXXXXXCTGAGTCGGAGACACGCA -3’, 10 µM) and H_2_O (total 50 μl) with the following PCR cycles: 98 °C for 30s, 7 cycles of 98 °C for 15 s, 55 °C for 30s, then 72°C for 30s, 72 °C for 2 min, 4 °C hold. For ATAC library amplification, 30 ng above product is mixed with 25 μl 2 × KAPA HiFi HotStart ReadyMix, 1.5 μl forward primer (Bioo-PCR-F) (10 µM), 1.5 μl reverse primer (Bioo-PCR-R: 5’-CAAGCAGAAGACGGCATACGAGAT-3’) and H_2_O (total 50 μl) with the following PCR cycles: 98 °C for 30s, 12 cycles of 98 °C for 10 s, 63 °C for 30s, 72°C for 1 min, then 72 °C for 2 mins, 4 °C hold. Enriched libraries were purified with 1.2 × volume of Ampure XP beads, and sequencing with Illumina Nextseq 2000 sequencer. For ATAC library, custom sequencing primers Read 1 (DEpi_ME_t1: 5’-GCCTCCCTCGCGCCATAGATGTGTATAAGAGACAG-3’), Read2 (DEpi_ME_t2: 5’-CTTGCCAGCCCGCTCAGAGATGTGTATAAGAGACAG-3’), Index 1 (Idx7: 5’ CTGTCTCTTATACACATCTCTGAGCGGGCTGGCAAG-3’), Index 2 (Idx5: 5’ CTGTCTCTTATACACATCTATGGCGCGAGGGAGGC-3’) were spiked-in during sequencing.

#### Preprocessing the wellDA-seq sequencing data

Based on the design of the cell barcode combinations in wellDA-seq, the BCL sequencing files were demultiplexed into FASTQ files for the DNA and ATAC modality separately by using bcl2fastq (v2.20.0.422) without allowing any mismatches of index sequences.

The FASTQ files of the DNA modality data were preprocessed by using a single cell DNA copy number calling pipeline described in a previous study^14^. Briefly, the FASTQ files were mapped to the human reference genome (hg19) with Bowtie2 (v2.5.3)^37^. Alignments with the equal start and end positions were marked as PCR duplicates, which were removed from processing. To estimate single-cell CNA profiles, the aligned reads were first counted across the genomic variable bins (about 220 kilobase per bin), resulting in bin counts.

Lowess Regression was then performed to normalize bin counts to calibrate GC content. The ratio of bin count to the average bin count of all bins in a cell was computed to represent the CNA ratio in each bin. Circular binary segmentation (‘alpha’ = 0.0001 and ‘undo.prune’ = 0.05) was performed on the bins using the package DNACopy (v1.68.0). The adjacent segments with non-significant differences in CNA ratios were merged by using function ‘mergeLevels’ of the package aCGH (v1.72.0), resulting in a single-cell segment ratios matrix. Any cell with high technical noise was excluded if it met any of the following criteria: 1) low average mapping quality (Q<1), 2) total read number was less than 100,000 when the average read number of all cells in the sample was about 1 million, and 3) there were greater than 10% of bins having no mapped reads.

The FASTQ files of the ATAC modality data were processed by using scATAC-pro (v1.4.1)^38^, which trims the reads using TrimGalore (v0.6.10) and maps the trimmed reads with Bowtie2 (v2.5.3). Alignments having a MAPQ score greater than 30 were used create the ATAC fragment files. We used the package Signac (v1.10.0) to preprocess the ATAC data. To determine the peaks of chromatin accessibility, we used the function ‘CallPeaks’ to read the fragments of all cells, which internally applies MACS2 (v2.2.7.1) with the parameters set as ‘--nomodel --shift -100 --extsize 200’ for peak calling. In the sample sequenced with multiple chips, all fragments were merged to call peaks. The singe-cell ATAC fragment count matrix was created by using the function ‘FeatureMatrix’. This matrix was binarized in the HBCA samples using ‘BinarizeCounts’ but was not binarized in the cancer cell line and DCIS data, thereby determining the single-cell chromatin accessibility profile. We further leveraged the package ArchR (v1.0.1)^39^ to calculate the transcription start sites (TSS) enrichment scores and the number of fragments for each cell. We kept cells having more than 1000 fragments and a TSS enrichment score of greater than 6 as previously described (https://www.encodeproject.org/atac-seq/). The cells passing the above quality control criteria for both DNA and ATAC modalities were used to downstream processing.

### Processing the ATAC modality data of wellDA-seq

The single-cell fragment count matrix was imported to Signac (v1.10.0)^40^ and used to run the function ‘RunTFIDF’ to perform normalization using Term Frequency-Inverse Document Frequency (TF-IDF). The peaks having the top total number of fragment counts were determined by ‘FindTopFeatures’ with the default parameters. Based on these peaks, we ran ‘RunSVD’ to perform linear dimensional reduction by singular value decomposition (SVD), which resulted in 100 latent semantic indexing (LSI) components. Any LSI component was disregarded in all the downstream analysis if it is highly correlated with the total fragments across cells (Pearson correlation coefficient > 0.75). Uniform Manifold Approximation and Projection (UMAP) was performed by running ‘RunUMAP’ to facilitate vislualization. Cell-to-cell similarity graph was built by running ‘FindNeighbors’ to facilitate downstream analysis of cell clustering.

### Inferring RNA expression from ATAC using Signac

The RNA expression of genes was inferred using the function ‘GeneActivity’ with the default parameters. This method counts the fragments overlapped with the upstream 2 kilobase genomic window of the TSS for each protein-coding gene in each single cell. Genomic locations and symbol names of the protein-coding genes were extracted from the database in the package ‘EnsDb.Hsapiens.v75’ (v2.99.0). The inferred gene activity matrix was normalized using the ‘NormalizedData’ with the parameter ‘scale.factor’ specified as the median of the total gene activity counts of cells.

### Identification of cell types using the ATAC modality data of wellDA-seq

To identify cell type, we determined the clusters of cells and then identified the top differentially expressed genes (DEGs) for each cluster. In details, we used the function ‘hdbscan’ of the package dbscan (v1.1-11) to perform cell clustering using the UMAP coordinates. To avoid under-/over-clustering, clustering was performed at a series of resolution with the parameter ‘minPts’ set from as fine as 5 to as coarse as 50 in an increasing step size of 5. We use the package ‘clustree’ (v0.4.4) to visualize the relationships between the clusters of using different clustering resolutions. To overcome over-clustering, we started from the smallest resolution leading to the maximum number of clusters and performed merging clusters. We determined the top DEGs for the clusters by applying the function ‘FindAllMarkers’ on the log-normalized inferred gene activity matrix with the default parameters. We only kept the DEGs that were detected in at least 1% of cells and had an FDR-adjusted p-value < 0.05 and an average log2-scaled fold change > 0. We merged clusters with similar top 50 DEGs while referring to the clustree’s result, which determined the final clusters. Based on the top DEGs of these clusters, we annotated the cell types for each cluster by referring to the known marker genes and transcription factors as previously reported^9,20^. A cluster showing less than 5 DEGs or noise-related genes were labelled as ‘noise’.

### Benchmarking wellDA-seq and single-cell DNA-seq techniques

To compare the technical performance of wellDA-seq versus the other four different scDNA-seq methods (Arc-well (p38 dataset), 10X genomics CNV, DLP+, and DOP-PCR), we first filtered out the cells with low-quality DNA copy number profiles from all five methods using a k-nearest neighbor (KNN) filtering method (resolution 0.8) implemented in CopyKit^41^ as described in previous study^14^. The overdispersion QC metric of genomic bin counts was calculated based on the following formula, in which Φ be the overdispersion parameter, μ be the mean read counts of genomic bins, and *D* be the index of dispersion as previously described^14,16^. To evaluate the breadth of coverage, the bam files were first downsampled to 750K to align the sequencing depth across all samples, then the Bedtools (v2.26.0)^42^ genomeCoverageBed function was applied to 120 randomly sampled single cells (to match with DOP-PCR dataset, which only had 122 cells left after QC) of each scDNA-seq method.

### Processing the DNA modality data of wellDA-seq

We leveraged Copykit to process the single-cell DNA data. The single-cell segment-ratio matrix was log_2_- scaled for normalization by using the function ‘logNorm’, resulting in a single-cell log-ratio matrix. Considering the ground state of genome copy number to be 2, we inferred the integer copy number by using the function ‘calcInteger’ which multiplies the segment-ratio matrix by 2.

### Identification of DNA subclones using the DNA modality data of wellDA-seq

To remove the doublets and noisy cells in the cancer cell line data, we applied the function ‘findOutliers’ with the parameter ‘resolution’=0.9 and ‘findAneuploidCells’ using the default parameters. However, in the samples of normal breast tissue and breast tumor, simply applying ‘findOutliers’ was likely to remove the rare cells harboring small CNA events. Therefore, we applied the function ‘findAneuploidCells’ with the parameter ‘resolution’ specified as ‘auto’ to split the cells into the tentative diploid and aneuploid cell groups, and further perform quality control and clustering analysis separately. In both cell groups, we performed principal component analysis (PCA) on the single-cell log_2_-ratio data for linear dimensional reduction using ‘runPca’.

In the tentative aneuploid cell group, we applied ‘runUmap’ with different ‘n_neighbors’ and ran ‘findClusters’ with different ‘k_superclones’ and ‘k_subclones’, depending on the total cell count N. If N < 200, ‘n_neighbors’ = 3, ‘k_superclones’ = 3, and ‘k_subclones’ = 3. If N is between 200 and 500, ‘n_neighbors’ = 5, ‘k_superclones’ = 5, and ‘k_subclones’ = 5. If N is between 500 and 1000, ‘n_neighbors’ = 10, ‘k_superclones’ = 10, and ‘k_subclones’ = 10. If N > 1000, ‘n_neighbors’ = 20, ‘k_superclones’ = 20, and ‘k_subclones’ = 20. To avoid over-clustering, subclones were merged if they had less than 3 consecutive bins with different CNA events. To identify the low-quality cells, we leveraged the function ‘calcInteger’ which internally uses Scquantum (v1.0.0) to compute two metrics ‘ploidy_score’ and ‘confidence_ratio’ for each cell. Cells having a ‘ploidy_score’ of smaller than 2 or a ‘confidence_ratio’ of greater than 0.05 were classified as low-quality or noisy cells, which were removed from downstream analysis. The remaining cells were classified as ‘aneuploid’.

In the tentative diploid cell group, we created the UMAP embedding by using ‘runUmap’ with the parameter ‘n_neighbors’=10. Then we identified the subclones using ‘findClusters’ with the parameter ‘k_superclones’=10 and ‘k_subclones’=10. To avoid over-clustering, subclones were merged if they had less than 3 consecutive bins with different CNA events. The subclones without any CNA events or having less than 10 consecutive CNA bins were classified as ‘diploid’. The remaining subclones were classified as ‘non-diploid’. If a ‘non-diploid’ subclone showed a high Pearson correlation coefficient with any ‘aneuploid’ subclone based on subclonal consensus CNA profiles, but it had an overall smaller log_2_-ratio value, this ‘non-diploid’ subclone was classified as a ‘doublet’ that was could be partly contaminated by a diploid cell. After removing the ‘doublet’ subclones, the remaining ‘non-diploid’ subclones were re-labelled as ‘aneuploid’.

### Processing of the combined DNA and ATAC modalities of wellDA-seq

After we processed the data from each modality separately, only cells that were remained in both CNA and ATAC modalities were kept for all downstream analysis.

### Copy number frequency in cancer cells

To calculate the frequency of the CNA events in cancer cells, we first established criteria for identifying can events based on log2 segment ratio values. Specifically, events with values greater than 1.3 or less than 0.7 were classified as CNAs. The frequency of these events was then calculated by dividing the number of cells with CNAs by the total number of cancer cells analyzed.

### Inferring chromatin hubs and RNA expression from ATAC using Cicero

We converted the Signac object containing the single-cell fragment count data to the object recognized by the package Monocle3 (v1.3.4)^43^ and Cicero (v1.3.9)^32^ by using the function ‘as_cell_data_set’, which allows us to perform the cis-co-accessible networks analysis to determine chromatin hubs, as described in the instructions (https://cole-trapnell-lab.github.io/cicero-release/docs_m3/#constructing-cis-regulatory-networks). Briefly, we ran the function ‘make_cicero_cds’ which aggregates similar cells to overcome the sparsity of single cell ATAC data, and then applied the function ‘run_cicero’ which estimates the peak-to-peak correlation. We focused on the positive correlated peak pairs representing the co-accessible peaks. Next, we sought the correlation threshold to remove the noisy co-accessible peaks. We sorted the peak pairs in an increasing order of the correlation values and identified the elbow point of the curve of the cumulative correlation values by using the function ‘find_curve_elbow’ of the package pathviewr (v1.1.7). The correlation value at this elbow point was used to set the parameter ‘coaccess_cutoff_override’ to run the function ‘generate_ccans’ which determines the chromatin hubs. This correlation value was also used in the ‘coaccess_cutoff’ parameter of the function ‘build_gene_activity_matrix’, which was applied to further infer the expression of the human hg19 genes based on the co-accessible peaks within the TSS ± 100kb windows. The human hg19 genes were extracted from the GTF file downloaded from Ensembl (https://grch37.ensembl.org/info/data/index.html).

### Inferring copy number alterations from the ATAC modality data

We utilized the same method to infer CNAs from ATAC as previously described^19^. The inferred CNAs were estimated in sliding genomic windows that are were in a step size of 2 megabase and each was 10 megabase wide. The inferred CNA score of each genomic window was the log_2_ fold change of signal compared with its 100 nearest neighbors in terms of the GC-content.

### Comparing the inferred CNAs with the directly measured CNAs of wellDA-seq

As the sliding windows used by the inferred CNAs and the genomic bins used by the measured CNAs of wellDA-seq have different genomic coordinates, we applied the function ‘findOverlaps’ of the package IRanges^44^ (v2.28.0) to map the sliding genomic windows to genomic bins. For each genomic bin, we computed the mean of the inferred CNA values of the overlapped sliding windows. This made the inferred CNAs extrapolated to the genomic bins and comparable to the CNAs results of wellDA-seq. For each subclone, we calculated the consensus CNAs using the function ‘calcConsensus’. Then we used the R function ‘cor.test’ to compute the Pearson correlation coefficient between the inferred and measured CNAs profiles and the p-value of correlation.

### Identifying subclonal genomic bins (SGBs)

In a comparison of two subclones, we used the integer copy number data and performed two-sided Wilcoxon test for each genomic bin by using the R function ‘wilcox.test’. P-values were adjusted by the Benjamini & Hochberg method via the R function ‘p.adjust’. SGBs were determined as the bins having an adjusted p-value smaller than 0.05 and a non-zero average integer copy number difference.

### Identifying differentially accessible peaks (DAPs)

Given sparsity of single-cell ATAC data, we first identified the informative peaks that were detected in at least 3 cells and showed a variance greater than the mean of the fragments in the cells of the two subclones in a comparison. For only each of these peaks, we performed a two-sided Wilcoxon test on the normalized fragment count data using the function ‘FindMarkers’ of Seurat which further corrects p-values by the Bonferroni method. DAPs were determined as the peaks having an adjusted p-value smaller than 0.05 and an average log_2_-fold change great than log_2_(1.1) (i.e., ∼0.14).

### Identify differentially accessible chromatin hubs (DACH)

We created a single-cell fragment count dataset for chromatin hubs by counting the fragments overlapping with the peaks designated to each chromatin hub in each cell. Then, we applied data normalization by using the function ‘RunTFIDF’. For each chromatin hub, we performed a two-sided Wilcoxon test on this normalized data using the function ‘FindMarkers’, which further corrects p-values using the Bonferroni method. DACHs were determined as the chromatin hubs having an adjusted p-value smaller than 0.05 and an average log_2_-fold change great than log_2_(1.1) (i.e., ∼0.14).

### Calculating the GtoE and EbyG percentages

The GtoE and EbyG percentages were computed based on the number of genomic bins of different categories. First, DACH-overlapped bins were determined as the genomic bins having at least 1bp overlaps with the peaks designated to DACHs. Then, DACH-overlapped SGBs were determined as the SGBs having at least 1bp overlaps with the peaks designated to DACHs. Finally, the GtoE percentage was computed as the ratio of the number of DACH-overlapped SGBs and SGBs, while the EbyG percentage was calculated as the ratio of the number of DACH-overlapped SGBs and DACH-overlapped genomic bins. Subclones with at least 15 cells were included in this analysis. If a subclone had less than 15 cells, but it comprised over 3% of cells in the sample, it was also included in this analysis.

### Module-based gene signature enrichment analysis (GSEA) for DACHs

To understand which gene expression was affected by DACHs, we applied the same 3-step module-based GSEA method as previously described^20^. Briefly, a collection of peaks designated to DACHs were termed as a module. Then we applied the function ‘AddChromatinModule’ of Signac to the ATAC fragment data to compute the module scores representing the overall enrichment of the DACHs in single cells. Next, we estimated the gene relevance by enumerating each gene and calculating the Pearson correlation coefficient between the module scores and the RNA expression values inferred by Cicero among single cells. Thus, the module obtained a gene-ranked vector that was decreasingly ordered by the coefficients. Finally, a regular GSEA was performed by using the function ‘GSEA’ of the package clusterProfiler (v4.2.2)^45^. Only the signatures having the Benjamini & Hochberg-adjusted p-value < 0.05 were considered significantly enriched.

### Inferring phylogenetic lineages using MEDICC2

Phylogenetic inference of consensus subclones was performed using MEDICC2^46^ based on the minimum event distances (medicc2 -a CN --total-copy-numbers). A diploid cell with copy number = 2 was added to the tree and designed as the root node, to root the tree. The integer copy number value of the first and the last 9 bins of each chromosome were assigned as the values of the first and the last 10^th^ bins to avoid chromosome border segmentation errors that could impact the tree construction. The trees were plotted by R package ‘ggtree’ (v3.2.1)^47^.

### Inferring segment size aware minimal evolution trees

To construct segment size aware tree in section “Genetic hardwiring and epigenetic plasticity in clonal lineages”, we computed the Manhattan distances between subclones using subclonal consensus integer copy number profiles. Then we used the function ‘fastme.bal’ of the package ape^48^ (v5.7-1) with the default parameters to reconstruct the minimal evolution tree.

### Calculating the global concordance of genotypes and chromatin accessibility profiles

If genotype and chromatin accessibility profile are highly concordant (or discordant), the subclones harboring the similar genotypes are expected to have similar (or distinct) chromatin accessibility profiles. To quantify the concordance level, we first computed the Manhattan distance using the subclonal consensus integer copy number profiles, representing the genetic distance between subclones. Then we computed the epigenetic subclone-to-subclone distances using the top 30 LSI components from the scATAC modality. To make the distances of the two modalities comparable, we used the function ‘rescale’ of the package scales (v1.2.1) to rescale this ATAC distance matrix to the value range of the CNAs distance matrix. Last, we used the R function ‘cor’ to compute the Pearson correlation coefficient of the subclone-to-subclone distances of the two modalities, which represented global concordance of genotypes and chromatin accessibility profiles in a sample.

### Estimating the overall expression of gene signatures

The overall expression of a gene signature was represented by the gene expression module score which was computed by applying the function ‘AddModuleScore_UCell’ of the package ‘UCell’ (v1.3.1) to the RNA expression data inferred by using Cicero.

### Calculating heritable and plastic gene signature scores

In each sample, we first tested if the module scores of a gene signature was significantly different across subclones by Kruskal-Wallis Rank Sum Test via the R function ‘kruskal.test’. We only focused on the gene signatures having a p-value < 0.05 in this analysis. To find the relative module scores of a gene signature across subclones, we computed the average UCell scores of the gene signature expressed in single cells of a subclone and then computed z-scores across subclones. Regarding samples, we excluded the breast cancer sample P5 from this analysis because there were no DACHs found in the supervised subclone-to-subclone comparisons. To quantify the extent to which a gene signature is heritable or plastic, we adopted the similar approach as computing the global concordance score. For each gene signature, we computed the Euclidean distances between subclones using the relative module scores. Finally, we estimated the correlation between this ATAC-based distances and the CNA-based distances by computing Pearson correlation coefficient. A higher (or low) correlation score indicated a heritable (or plastic) gene signature.

## References

1. Łukasiewicz, S. et al. Breast Cancer-Epidemiology, Risk Factors, Classification, Prognostic Markers, and Current Treatment Strategies-An Updated Review. Cancers (Basel) 13, 4287 (2021).

2. Martelotto, L. G., Ng, C. K., Piscuoglio, S., Weigelt, B. & Reis-Filho, J. S. Breast cancer intra-tumor heterogeneity. Breast Cancer Research 16, 210 (2014).

3. Turashvili, G. & Brogi, E. Tumor Heterogeneity in Breast Cancer. Front. Med. 4, (2017).

4. Shen, H. & Laird, P. W. Interplay between the Cancer Genome and Epigenome. Cell 153, 38–55 (2013).

5. Corces, M. R. et al. The chromatin accessibility landscape of primary human cancers. Science 362, eaav1898 (2018).

6. Hoadley, K. A. et al. Cell-of-Origin Patterns Dominate the Molecular Classification of 10,000 Tumors from 33 Types of Cancer. Cell 173, 291–304.e6 (2018).

7. Marusyk, A., Janiszewska, M. & Polyak, K. Intratumor Heterogeneity: The Rosetta Stone of Therapy Resistance. Cancer Cell 37, 471–484 (2020).

8. Orrantia-Borunda, E., Anchondo-Nuñez, P., Acuña-Aguilar, L. E., Gómez-Valles, F. O. & Ramírez-Valdespino, C. A. Subtypes of Breast Cancer. in Breast Cancer (ed. Mayrovitz, H. N.) (Exon Publications, Brisbane (AU), 2022).

9. Kumar, T. et al. A spatially resolved single-cell genomic atlas of the adult human breast. Nature 620, 181–191 (2023).

10. Nguyen, Q. H. et al. Profiling human breast epithelial cells using single cell RNA sequencing identifies cell diversity. Nature Communications 9, 2028 (2018).

11. Lim, E. et al. Aberrant luminal progenitors as the candidate target population for basal tumor development in BRCA1 mutation carriers. Nat Med 15, 907–913 (2009).

12. Molyneux, G., et al. *BRCA1* Basal-like Breast Cancers Originate from Luminal Epithelial Progenitors and Not from Basal Stem Cells. Cell Stem Cell 7, 403–417 (2010).

13. Bhat-Nakshatri, P., et al. A single-cell atlas of the healthy breast tissues reveals clinically relevant clusters of breast epithelial cells. CR Med 2, (2021).

14. Wang, K. et al. Archival single-cell genomics reveals persistent subclones during DCIS progression. Cell 186, 3968–3982.e15 (2023).

15. Laks, E. et al. Clonal Decomposition and DNA Replication States Defined by Scaled Single-Cell Genome Sequencing. Cell 179, 1207–1221.e22 (2019).

16. Minussi, D. C. et al. Breast tumours maintain a reservoir of subclonal diversity during expansion. Nature 592, 302–308 (2021).

17. Cusanovich, D. A. et al. Multiplex single-cell profiling of chromatin accessibility by combinatorial cellular indexing. Science 348, 910–914 (2015).

18. Lareau, C. A. et al. Droplet-based combinatorial indexing for massive-scale single-cell chromatin accessibility. Nature Biotechnology 37, 916–924 (2019).

19. Satpathy, A. T. et al. Massively parallel single-cell chromatin landscapes of human immune cell development and intratumoral T cell exhaustion. Nature Biotechnology 37, 925–936 (2019).

20. Wang, K. et al. Simple oligonucleotide-based multiplexing of single-cell chromatin accessibility. Molecular Cell 81, 4319–4332.e10 (2021).

21. Nikolic, A. et al. Copy-scAT: Deconvoluting single-cell chromatin accessibility of genetic subclones in cancer. Science Advances 7, eabg6045 (2021).

22. Wu, C.-Y. et al. Integrative single-cell analysis of allele-specific copy number alterations and chromatin accessibility in cancer. Nat Biotechnol 39, 1259–1269 (2021).

23. Ramakrishnan, A. et al. epiAneufinder identifies copy number alterations from single-cell ATAC-seq data. Nat Commun 14, 5846 (2023).

24. Tedesco, M. et al. Chromatin Velocity reveals epigenetic dynamics by single-cell profiling of heterochromatin and euchromatin. Nat Biotechnol 40, 235–244 (2022).

25. Queitsch, K. et al. Accessible high-throughput single-cell whole-genome sequencing with paired chromatin accessibility. Cell Reports Methods 3, 100625 (2023).

26. Gao, R. et al. Punctuated copy number evolution and clonal stasis in triple-negative breast cancer. Nature Genetics 48, 1119–1130 (2016).

27. Yizhak, K. et al. RNA sequence analysis reveals macroscopic somatic clonal expansion across normal tissues. Science 364, eaaw0726 (2019).

28. Martincorena, I. et al. Somatic mutant clones colonize the human esophagus with age. Science 362, 911–917 (2018).

29. Jakubek, Y. A. et al. Large-scale analysis of acquired chromosomal alterations in non-tumor samples from patients with cancer. Nat Biotechnol 38, 90–96 (2020).

30. Li, R. et al. A body map of somatic mutagenesis in morphologically normal human tissues. Nature 597, 398–403 (2021).

31. Zhou, Y. et al. Single-Cell Multiomics Sequencing Reveals Prevalent Genomic Alterations in Tumor Stromal Cells of Human Colorectal Cancer. Cancer Cell 38, 818–828.e5 (2020).

32. Pliner, H. A. et al. Cicero Predicts *cis*-Regulatory DNA Interactions from Single-Cell Chromatin Accessibility Data. Molecular Cell 71, 858–871.e8 (2018).

33. Liberzon, A. et al. Molecular signatures database (MSigDB) 3.0. Bioinformatics 27, 1739–1740 (2011).

34. Cancer Genome Atlas Network. Comprehensive molecular portraits of human breast tumours. Nature 490, 61–70 (2012).

35. Patel, A. P. et al. Single-cell RNA-seq highlights intratumoral heterogeneity in primary glioblastoma. Science 344, 1396–1401 (2014).

36. Tyler, M. & Tirosh, I. Decoupling epithelial-mesenchymal transitions from stromal profiles by integrative expression analysis. Nat Commun 12, 2592 (2021).

37. Langmead, B. & Salzberg, S. L. Fast gapped-read alignment with Bowtie 2. Nat Methods 9, 357–359 (2012).

38. Yu, W., Uzun, Y., Zhu, Q., Chen, C. & Tan, K. scATAC-pro: a comprehensive workbench for single-cell chromatin accessibility sequencing data. Genome Biology 21, 94 (2020).

39. Granja, J. M. et al. ArchR is a scalable software package for integrative single-cell chromatin accessibility analysis. Nat Genet 53, 403–411 (2021).

40. Stuart, T., Srivastava, A., Madad, S., Lareau, C. A. & Satija, R. Single-cell chromatin state analysis with Signac. Nat Methods 18, 1333–1341 (2021).

41. Minussi, D. C. et al. Resolving clonal substructure from single cell genomic data using CopyKit. 2022.03.09.483497 Preprint at 10.1101/2022.03.09.483497 (2022).

42. Quinlan, A. R. & Hall, I. M. BEDTools: a flexible suite of utilities for comparing genomic features. Bioinformatics 26, 841–842 (2010).

43. Qiu, X. et al. Reversed graph embedding resolves complex single-cell trajectories. Nat Methods 14, 979–982 (2017).

44. Lawrence, M. et al. Software for Computing and Annotating Genomic Ranges. PLOS Computational Biology 9, e1003118 (2013).

45. Yu, G., Wang, L.-G., Han, Y. & He, Q.-Y. clusterProfiler: an R Package for Comparing Biological Themes Among Gene Clusters. OMICS: A Journal of Integrative Biology 16, 284–287 (2012).

46. Kaufmann, T. L. et al. MEDICC2: whole-genome doubling aware copy-number phylogenies for cancer evolution. Genome Biology 23, 241 (2022).

47. Yu, G., Smith, D. K., Zhu, H., Guan, Y. & Lam, T. T.-Y. ggtree: an r package for visualization and annotation of phylogenetic trees with their covariates and other associated data. Methods in Ecology and Evolution 8, 28–36 (2017).

48. Paradis, E., Claude, J. & Strimmer, K. APE: Analyses of Phylogenetics and Evolution in R language. Bioinformatics 20, 289–290 (2004).

