## supplementary figures and tables for "Single cell genome and epigenome co-profiling reveals hardwiring and plasticity in breast cancer"

**a** ATAC library sequence structure

ATAC tagmentation (Related to I. Cell suspension tagmentation in main Figure 1a)

```
5'-GCCTCCCTCGCGCCATAGATGTGTATAAGAGACAG------------DNA--------   pCTGTCTCTTATACACATCT-3'
3'                  TCTACACATATTCTCTGTCp    --------DNA-----------GACAGAGAATATGTGTAGAGACTCGCCCGACCGTTC-5'
```

Gap filling (Related to III. Single cell DNA & ATAC dual indexing: 3. PCR and cell barcoding)

```
5'-GCCTCCCTCGCGCCATAGATGTGTATAAGAGACAG------------DNA-----------CTGTCTCTTATACACATCTCTGAGCGGGCTGGCAAG-3'
3'-CGGAGGGAGCGCGGTATCTACACATATTCTCTGTC------------DNA-----------GACAGAGAATATGTGTAGAGACTCGCCCGACCGTTC-5'
```

DNA&ATAC co-enrichment PCR (Related to III. Single cell DNA & ATAC dual indexing: 3. PCR and cell barcoding)

PCR Primers (Table S5):

5'-AATGATACGGCGACCACCGAGATCTACACTGA-**ATAC_CB1**-GCCTCCCTCGCGCCAT-3'
5'-CAAGCAGAAGACGGCATACGAGAT-**ATAC_CB2**-CTTGCCAGCCCGCTCAG-3'

```
5'-AATGATACGGCGACCACCGAGATCTACACTGA-ATAC_CB1-GCCTCCCTCGCGCCATAGATGTGTATAAGAGACAG-------DNA-------
3'-TTACTATGCCGCTGGTGGCTCTAGATGTGACT-ATAC_CB1-CGGAGGGAGCGCGGTATCTACACATATTCTCTGTC-------DNA-----
---CTGTCTCTTATACACATCTCTGAGCGGGCTGGCAAG-ATAC_CB2-ATCACGATCTCGTATGCCGTCTTCTGCTTG-3'
----- GACAGAGAATATGTGTAGAGACTCGCCCGACCGTTC-ATAC_CB2-TAGTGCTAGAGCATACGGCAGAAGACGAAC-5'
```

ATAC enrichment PCR (Related to V. Modality enrichment PCR)

PCR Primers:

5'-AATGATACGGCGACCACCGAGATCTACAC-3'
5'-CAAGCAGAAGACGGCATACGAGAT-3'

```
5'-AATGATACGGCGACCACCGAGATCTACACTGA-ATAC_CB1-GCCTCCCTCGCGCCATAGATGTGTATAAGAGACAG-------DNA-------
3'-TTACTATGCCGCTGGTGGCTCTAGATGTGACT-ATAC_CB1-CGGAGGGAGCGCGGTATCTACACATATTCTCTGTC-------DNA-----
---CTGTCTCTTATACACATCTCTGAGCGGGCTGGCAAG-ATAC_CB2-ATCACGATCTCGTATGCCGTCTTCTGCTTG-3'
-----GACAGAGAATATGTGTAGAGACTCGCCCGACCGTTC-ATAC_CB2-TAGTGCTAGAGCATACGGCAGAAGACGAAC-5'
```

**ATAC_CB1 / ATAC_CB2**: 8bp ATAC cell barcodes

**b** DNA library sequence structure

DNA tagmentation (Related to III. Single cell DNA & ATAC dual indexing: 2. Tagmentation)

```
5'-TCGTCGGCAGCGTCAGATGTGTATAAGAGACAG------------DNA--------   pCTGTCTCTTATACACATCT-3'
3'                TCTACACATATTCTCTGTCp    --------DNA-----------GACAGAGAATATGTGTAGAGGCTCGGGTGCTCTG-5'
```

Gap filling (Related to III. Single cell DNA & ATAC dual indexing: 3. PCR and cell barcoding)

```
5'-TCGTCGGCAGCGTCAGATGTGTATAAGAGACAG------------DNA-----------CTGTCTCTTATACACATCTCCGAGCCCACGAGAC-3'
3'-AGCAGCCGTCGCAGTCTACACATATTCTCTGTC------------DNA-----------GACAGAGAATATGTGTAGAGGCTCGGGTGCTCTG-5'
```

DNA&ATAC co-enrichment PCR (Related to III. Single cell DNA & ATAC dual indexing: 3. PCR and cell barcoding)

PCR Primers (Table S6):

5'-AATGATACGGCGACCACCGAGATCTACAC-**DNA_CB1**-TCGTCGGCAGCGTC-3'
5'-CTGAGTCGGAGACACGCA-**DNA_CB2**-GTCTCGTGGGCTCGG-3'

```
5'-AATGATACGGCGACCACCGAGATCTACAC-DNA_CB1-TCGTCGGCAGCGTCAGATGTGTATAAGAGACAG-------DNA-----
3'-TTACTATGCCGCTGGTGGCTCTAGATGTG-DNA_CB1-AGCAGCCGTCGCAGTCTACACATATTCTCTGTC-------DNA----
-CTGTCTCTTATACACATCTCCGAGCCCACGAGAC-DNA_CB2-TGCGTGTCTCCGACTCAG-3'
--GACAGAGAATATGTGTAGAGGCTCGGGTGCTCTG-DNA_CB2-ACGCACAGAGGCTGAGTC-5'
```

DNA enrichment PCR (Related to V. Modality enrichment PCR)

PCR Primers:

5'-AATGATACGGCGACCACCGAGATCTACAC-3'
5'-CAAGCAGAAGACGGCATACGAGAT-**DNA_CB3**-CTGAGTCGGAGACACGCA-3'

```
5'-AATGATACGGCGACCACCGAGATCTACAC-DNA_CB1-TCGTCGGCAGCGTCAGATGTGTATAAGAGACAG-------DNA-- ------
3'-TTACTATGCCGCTGGTGGCTCTAGATGTG-DNA_CB1-AGCAGCCGTCGCAGTCTACACATATTCTCTGTC-------DNA-- ---
---CTGTCTCTTATACACATCTCCGAGCCCACGAGAC-DNA_CB2-TGCGTGTCTCCGACTCAG-DNA_CB3-ATCTCGTATGCCGTCTTCTGCTTG-3'
------ GACAGAGAATATGTGTAGAGGCTCGGGTGCTCTG-DNA_CB2-ACGCACAGAGGCTGAGTC-DNA_CB3-TAGAGCATACGGCAGAAGACGAAC-5'
```

**DNA_CB1 / DNA_CB2 / DNA_CB3**: 8bp DNA cell barcodes

### Extended Data Fig. 1 Detailed wellDA-seq DNA and ATAC library sequence structure

**a.** The wellDA-seq ATAC library sequence structure after different reaction steps. **b.** The wellDA-seq DNA library sequence structure after different reaction steps.

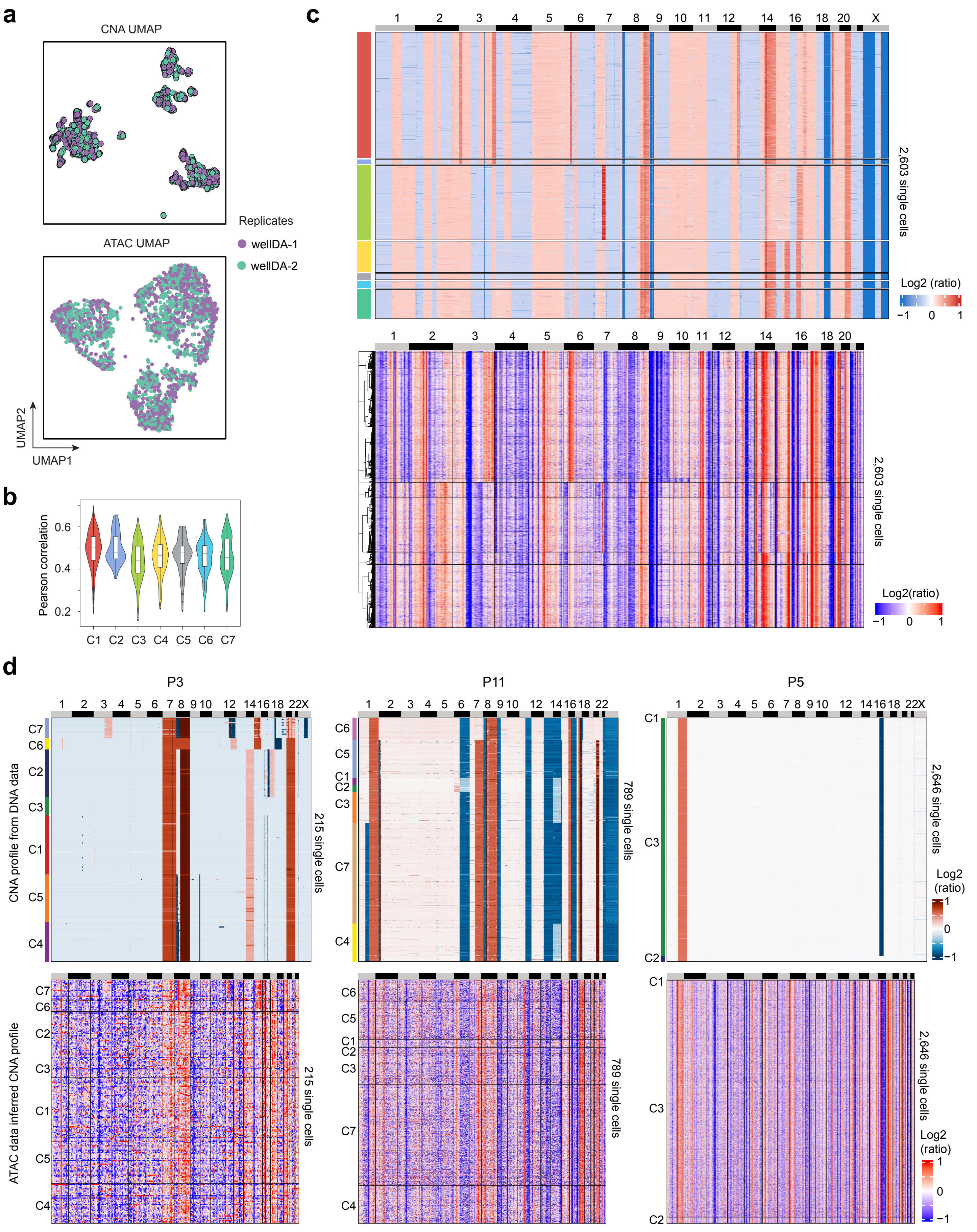
a
CNA UMAP
ATAC UMAP
Replicates
wellDA-1
wellDA-2
UMAP2
UMAP1
b
Pearson correlation
0.6
0.4
0.2
C1 C2 C3 C4 C5 C6 C7
c
1 2 3 4 5 6 7 8 9 10 11 12 14 16 18 20 X
2,603 single cells
Log2 (ratio)
−1 0 1
1 2 3 4 5 6 7 8 9 10 11 12 14 16 18 20
2,603 single cells
Log2(ratio)
−1 0 1
d
P3
P11
P5
CNA profile from DNA data
ATAC data inferred CNA profile
215 single cells
789 single cells
2,646 single cells
C1
C2
C3
C4
C5
C6
C7
Log2 (ratio)
1
0
−1

**Extended Data Fig. 2 Comparing actual DNA copy number profiles to ATAC-inferred CNA profiles**

**a.** Batch effect evaluation of two replicate wellDA-seq experiments from the MDA231 cell line. UMAPs of single cell DNA copy number profiles (top panel) and single cell ATAC profiles (bottom panel) from two wellDA-seq experiments. **b.** Pearson correlation of single cell CNA profiles of each DNA subclone identified using the DNA data from wellDA-seq and the inferred single cell CNA profiles from the ATAC data. **c.** Heatmaps of DNA copy number profiles of 2,603 single cells using the DNA modality data (top panel) of wellDA-seq and the inferred single cell copy number profiles of the same 2,603 single cells using the ATAC modality data (bottom panel) of wellDA-seq. **d.** Heatmaps of single cell DNA copy number profiles from the wellDA-seq DNA modality data (top panel) compared to the inferred single cell copy number profiles from ATAC modality data (bottom panel). Data were generated from three different breast tumors using wellDA-seq.

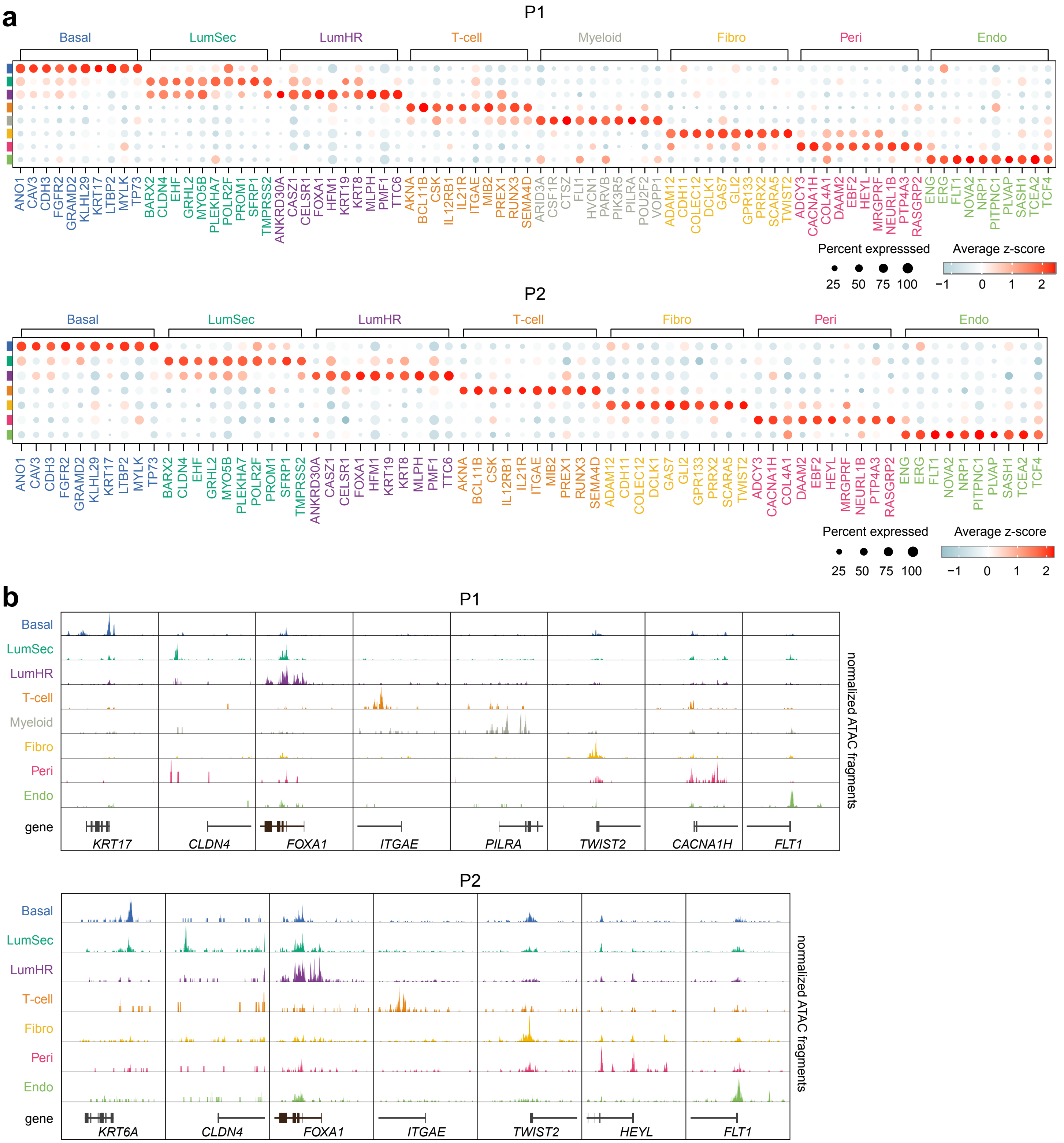

**Extended Data Fig. 3 Identifying cell types in breast tissue samples from two healthy women**

**a.** Dotplots of P1 and P2 showing the relative RNA expression inferred from ATAC profiles of 10 top differential expressed genes in each cell type. **b.** Track plots of P1 and P2 showing the normalized ATAC fragment counts of cell types in the 200kb genomic window surrounding transcriptional start sites of the cell type-specific genes.

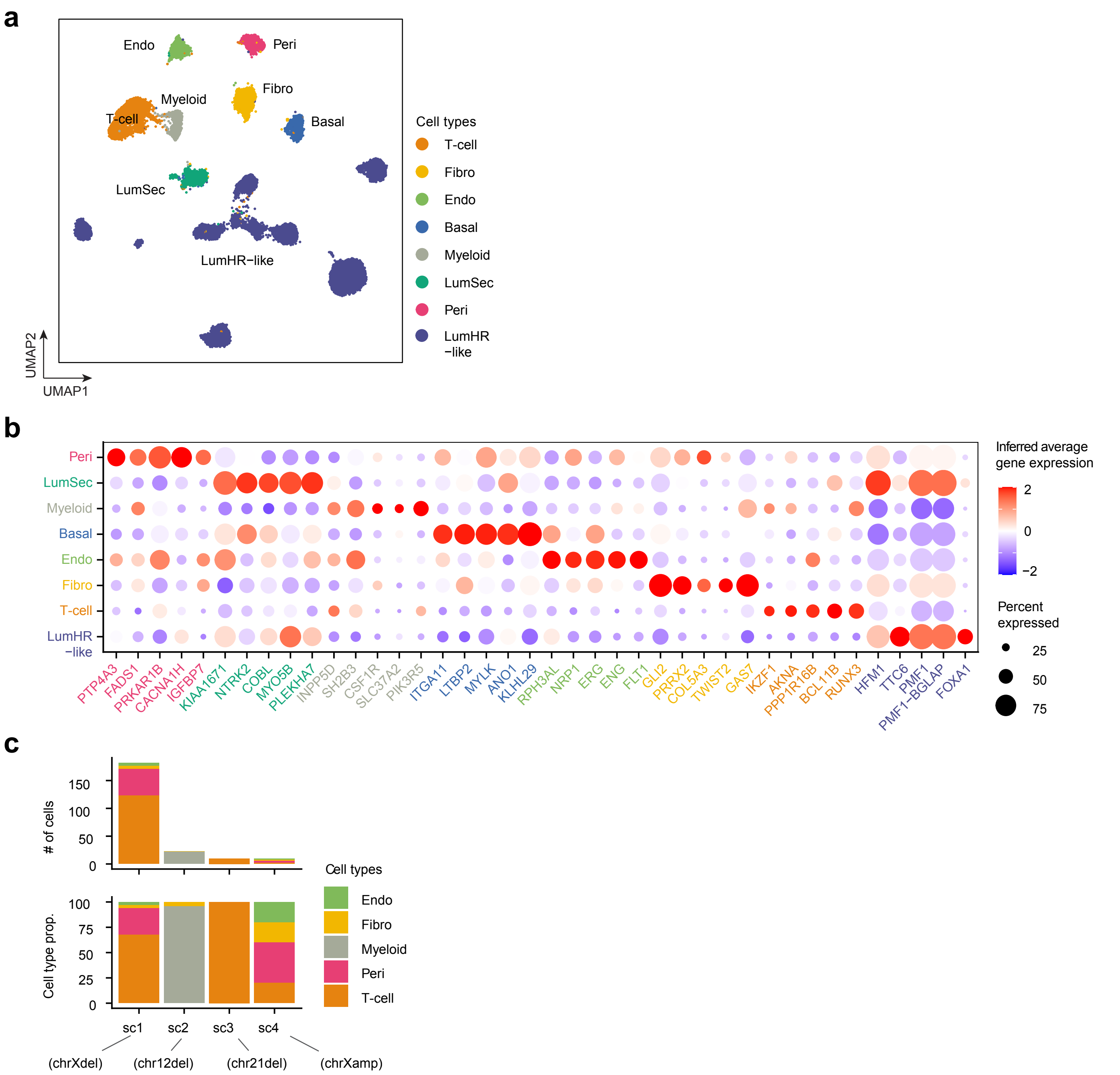

**Extended Data Fig. 4 Cell type identities and top marker genes of 9 breast cancers**

**a.** ATAC UMAP showing 8 major cell types in 9 breast cancers identified by wellDA-seq. **b.** Heatmap of top 5 marker genes from each cell type cluster from a healthy woman (P1) are used to annotate the cell type clusters across 9 merged breast cancers. **c.** Number of cells and cell type proportions of 4 different non-epithelial aneuploid subclones.

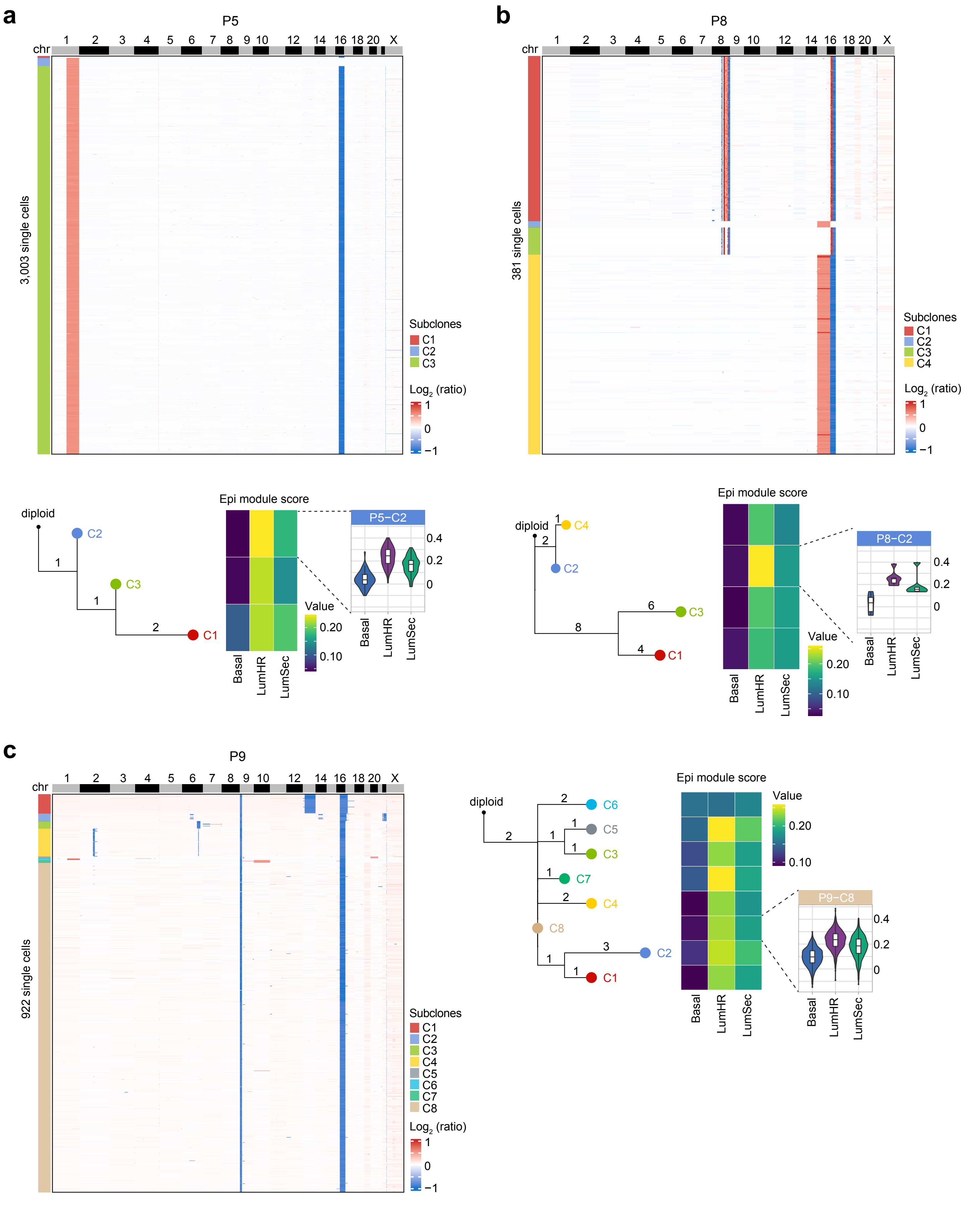
a
P5
chr
1 2 3 4 5 6 7 8 9 10 12 14 16 18 20 X
3,003 single cells
Subclones
C1
C2
C3
Log2 (ratio)
1
0
−1
diploid
C2
1
C3
1
2
C1
Epi module score
P5−C2
0.4
0.2
0
Value
0.20
0.10
Basal
LumHR
LumSec
b
P8
chr
1 2 3 4 5 6 7 8 9 10 12 14 16 18 20 X
381 single cells
Subclones
C1
C2
C3
C4
Log2 (ratio)
1
0
−1
diploid
1
C4
2
C2
6
C3
8
4
C1
Epi module score
P8−C2
0.4
0.2
0
Value
0.20
0.10
Basal
LumHR
LumSec
c
P9
chr
1 2 3 4 5 6 7 8 9 10 12 14 16 18 20 X
922 single cells
Subclones
C1
C2
C3
C4
C5
C6
C7
C8
Log2 (ratio)
1
0
−1
diploid
2
C6
1
C5
2
1
1
C3
1
C7
2
C4
C8
3
C2
1
1
C1
Epi module score
Value
0.20
0.10
P9−C8
0.4
0.2
0
Basal
LumHR
LumSec

**Extended Data Fig. 5 Additional samples showing epithelial identities of ancestral cancer cells**

**a.** Clustered heatmap of single cell copy number profiles of cancer cells, with phylogenetic tree of subclonal consensus integer copy number profiles and the epithelial cell type module score of each cancer subclone from patients P5, P8 (**b**) and P9 (**c**).

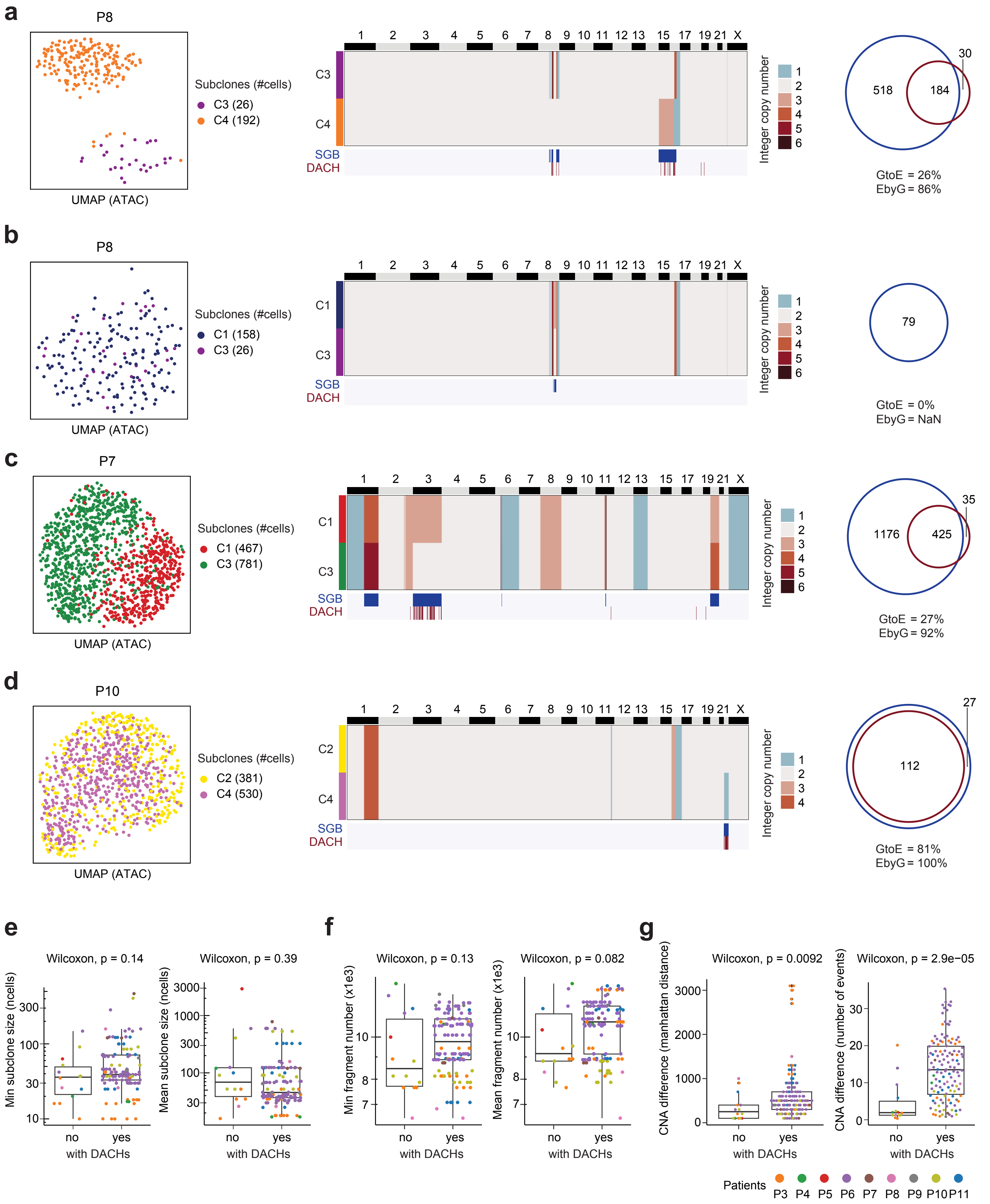
a
P8
Subclones (#cells)
C3 (26)
C4 (192)
UMAP (ATAC)
1 2 3 4 5 6 7 8 9 10 11 12 13 15 17 19 21 X
C3
C4
SGB
DACH
Integer copy number
1 2 3 4 5 6
518
184
30
GtoE = 26%
EbyG = 86%
b
P8
Subclones (#cells)
C1 (158)
C3 (26)
UMAP (ATAC)
C1
C3
SGB
DACH
Integer copy number
79
GtoE = 0%
EbyG = NaN
c
P7
Subclones (#cells)
C1 (467)
C3 (781)
UMAP (ATAC)
C1
C3
SGB
DACH
Integer copy number
1176
425
35
GtoE = 27%
EbyG = 92%
d
P10
Subclones (#cells)
C2 (381)
C4 (530)
UMAP (ATAC)
C2
C4
SGB
DACH
Integer copy number
1 2 3 4
112
27
GtoE = 81%
EbyG = 100%
e
Wilcoxon, p = 0.14
Min subclone size (ncells)
Wilcoxon, p = 0.39
Mean subclone size (ncells)
no yes
with DACHs
f
Wilcoxon, p = 0.13
Min fragment number (x1e3)
Wilcoxon, p = 0.082
Mean fragment number (x1e3)
g
Wilcoxon, p = 0.0092
CNA difference (manhattan distance)
Wilcoxon, p = 2.9e−05
CNA difference (number of events)
Patients
P3 P4 P5 P6 P7 P8 P9 P10 P11

**Extended Data Fig. 6 Additional samples showing impact of subclonal CNAs on chromatin accessibility**

**a-d**, Examples of the subclone comparisons of DNA and ATAC modalities in three different patients (P8, P7 and P10). Panels are displayed from left to right showing the UMAP of the ATAC modality data colored by DNA subclones, the heatmap of subclone consensus copy number profiles of two compared subclones with SGB and DACH events annotated, and Venn diagram of the overlapping of the SGBs and DACHs of the two compared subclones C3 and C4 in P8 (**a**), C1 and C3 in P8 (**b**), C1 and C3 in P7 (**c**), and C2 and C4 in P10 (**d**). NaN: 'not a number' because of 0/0 values. **e,f,** Boxplot showing the minimum (left panel) and average (right panel) of subclonal size (**e**) and ATAC fragment counts (**f**) in each subclone-to-subclone comparison (dots) grouped by the presence of DACHs. **g,** Boxplot showing the CNA distance based on the CNA events (left panel) and Manhattan distance (right panel) in each subclone-to-subclone comparison (dots) grouped by the existence of DACHs. P-values were obtained from Wilcoxon signed-rank test. In **e, f,** y-axis is shown on a $\log_{10}$ scale.

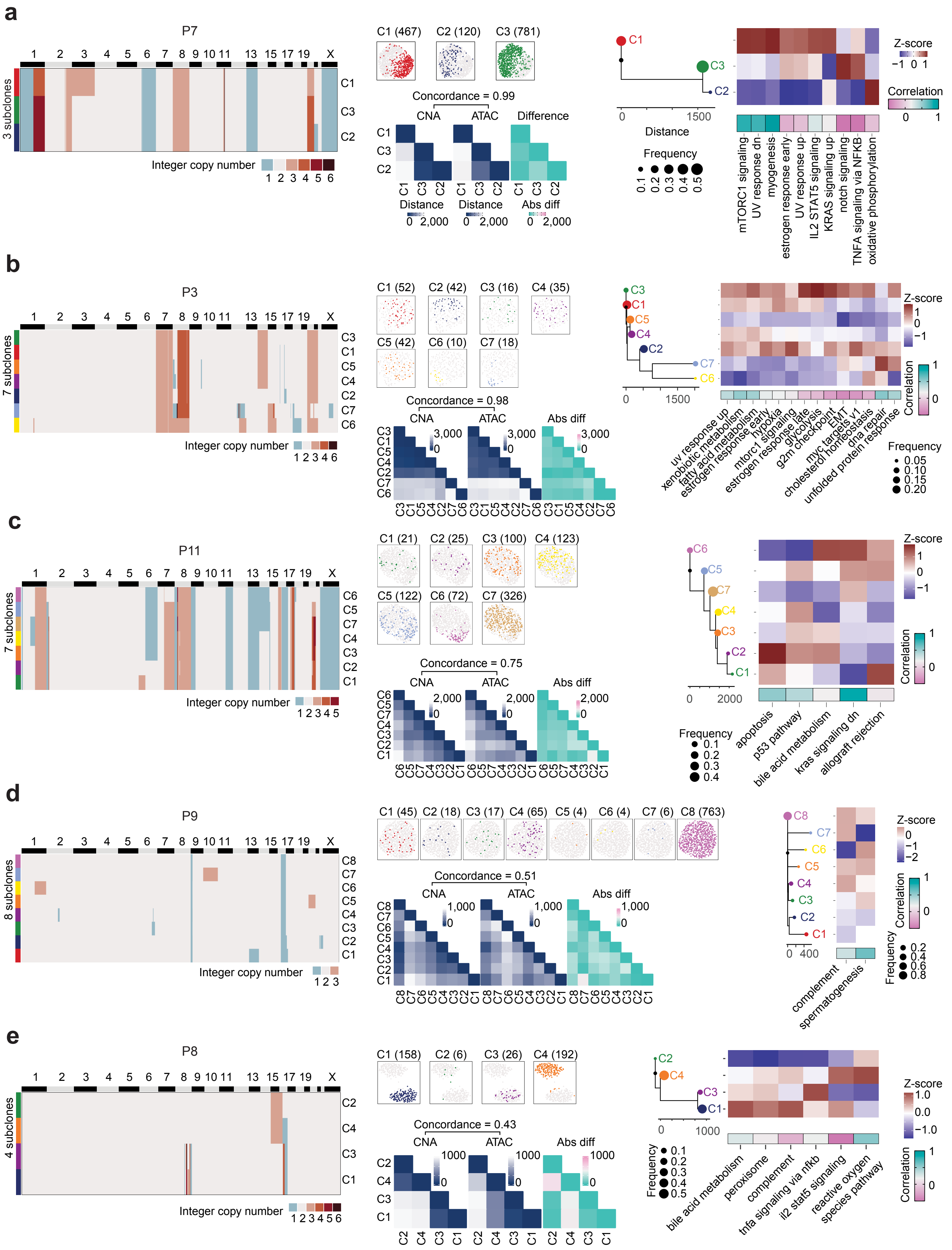
a
P7
3 subclones
C1
C3
C2
Integer copy number
1 2 3 4 5 6
C1 (467) C2 (120) C3 (781)
Concordance = 0.99
CNA
ATAC
Difference
Distance
Distance
Abs diff
0 2,000
Distance
1500
Frequency
0.1 0.2 0.3 0.4 0.5
Z-score
−1 0 1
Correlation
0 1
mTORC1 signaling
UV response dn
myogenesis
estrogen response early
UV response up
IL2 STAT5 signaling
KRAS signaling up
notch signaling
TNFA signaling via NFKB
oxidative phosphorylation
b
P3
7 subclones
C3
C1
C5
C4
C2
C7
C6
Integer copy number
1 2 3 4 6
C1 (52) C2 (42) C3 (16) C4 (35)
C5 (42) C6 (10) C7 (18)
Concordance = 0.98
CNA
ATAC
Abs diff
3,000
0
1500
Z-score
Correlation
Frequency
0.05
0.10
0.15
0.20
uv response up
xenobiotic metabolism
fatty acid metabolism
estrogen response early
hypoxia
mtorc1 signaling
estrogen response late
glycolysis
g2m checkpoint
myc targets v1
EMT
cholesterol homeostasis
dna repair
unfolded protein response
c
P11
7 subclones
C6
C5
C7
C4
C3
C2
C1
Integer copy number
1 2 3 4 5
C1 (21) C2 (25) C3 (100) C4 (123)
C5 (122) C6 (72) C7 (326)
Concordance = 0.75
CNA
ATAC
Abs diff
2,000
0
2000
Z-score
Correlation
Frequency
0.1
0.2
0.3
0.4
apoptosis
p53 pathway
bile acid metabolism
kras signaling dn
allograft rejection
d
P9
8 subclones
C8
C7
C6
C5
C4
C3
C2
C1
Integer copy number
1 2 3
C1 (45) C2 (18) C3 (17) C4 (65) C5 (4) C6 (4) C7 (6) C8 (763)
Concordance = 0.51
CNA
ATAC
Abs diff
1,000
0
400
Z-score
0
−1
−2
Correlation
Frequency
0.2
0.4
0.6
0.8
complement
spermatogenesis
e
P8
4 subclones
C2
C4
C3
C1
Integer copy number
1 2 3 4 5 6
C1 (158) C2 (6) C3 (26) C4 (192)
Concordance = 0.43
CNA
ATAC
Abs diff
1000
0
Frequency
0.1
0.2
0.3
0.4
0.5
Z-score
1.0
0.0
−1.0
Correlation
bile acid metabolism
peroxisome
complement
tnfa signaling via nfkb
il2 stat5 signaling
reactive oxygen species pathway

**Extended Data Fig. 7 Genetic hardwiring and epigenetic plasticity in 5 additional breast cancer samples**

**a-e,** Panels from left to right show the heatmap of the consensus integer CNA profiles of subclones, the UMAP plot of ATAC modality data colored by subclones, heatmaps (middle-lower panel) of the subclone-to-subclone distance matrix based on CNA, ATAC, and the difference of these two matrices, the minimum evolution tree, the heatmap of the relative gene activities of the gene signatures (top right) and the heritability/plasticity scores (bottom right) using the wellDA-seq data of the cancer cells in the breast tumor samples from P7 (**a**), P3 (**b**), P11 (**c**), P9 (**d**) and P8 (**e**).

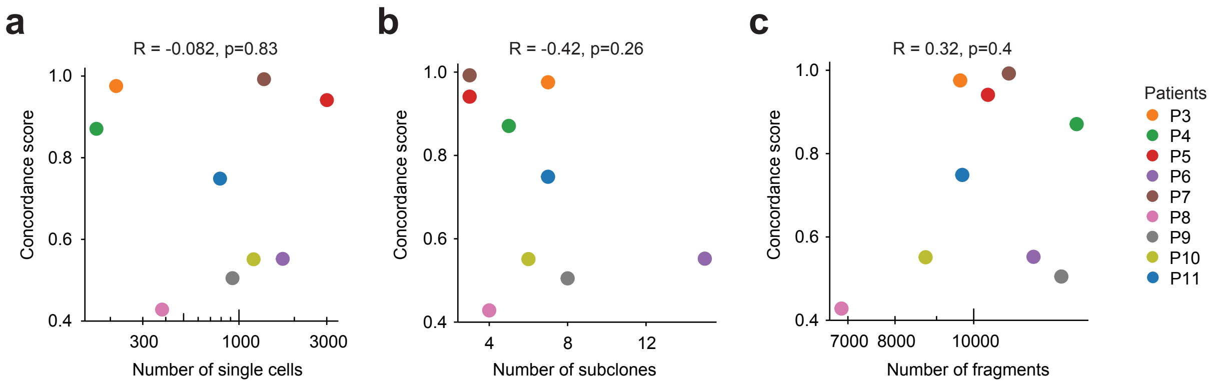

**Extended Data Fig. 8 Assessing the CNA-ATAC concordance score with technical factors**

**a-c,** Calculating the Pearson correlation coefficient and p-values between the CNA-ATAC concordance and the cell count (**a**), the subclone count (**b**), and the average fragment number (**c**). In **a,c**, the x-axis was shown in a $log_{10}$ scale.

| Clinical information of 2 healthy women and 9 patients with breast cancers | | | | | | | | | |
|---|---|---|---|---|---|---|---|---|---|
| Individual/ patient | Sample_ID | ER status | PR status | HER2 status | Grade | Histology | pT | pN | Age range |
| P1 | BCMHBCA52R | NA | NA | NA | NA | NA | NA | NA | 25-30 |
| P2 | BCMHBCA66L | NA | NA | NA | NA | NA | NA | NA | 40-45 |
| P3 | BCMDCIS22T | Pos | Pos | NA | G3, poorly differentiated | IDC | pT2 | pNX | 45-50 |
| P4 | BCMDCIS28T | Pos | Pos | Neg | G2, moderately differentiated | IDC | pT2 | pN1 | 50-55 |
| P5 | BCMDCI35T | Pos | Pos | Neg | G1, well differentiated | IDC | mpT1a | mpN0 | 60-65 |
| P6 | BCMDCIS41T | Pos | Pos | NA | DCIS only | DCIS only | pTis (DCIS) | pNx | 50-55 |
| P7 | BCMDCIS51T | Pos | Pos | Neg | G3, poorly differentiated | IDC | pT2 | snpN1a | 60-65 |
| P8 | BCMDCIS66T | Pos | Pos | NA | DCIS only; G2 | DCIS only | pTis (DCIS) | pNx | 40-42 |
| P9 | BCMDCIS67T | Pos | Pos | NA | DCIS only; G2 | DCIS only | pTis (DCIS) | pN0 | 65-70 |
| P10 | BCMDCIS68T | Pos | Pos | Neg | G2, moderately differentiated | IDC | pT1b | pNx | 75-80 |
| P11 | BCMDCIS69T | Pos | Pos | Neg | G3, poorly differentiated | IDC | pT1c | pN1 | 55-60 |

Pos: Positive

Neg: Negative

NA: Not Available

**Supplementary Table 1 Clinical metadata for 2 healthy women and 9 breast cancer patients**

Clinical information of the 2 normal breast tissues from disease-free women and 9 breast cancers that were analyzed by wellDA-seq in this study. ER and PR positive status was defined by Immunohistochemistry (IHC) with greater than 1% staining, while HER2 positive status was defined by fluorescence in situ hybridization (FISH) with scores of 2+ or 3+. Histopathological analysis of the tissue was used to classify the lesions as ductal carcinoma in situ (DCIS) or invasive ductal carcinoma (IDC) with DCIS and their tumor grades. The clinical staging was determined by the tumor size (pT) and nodal involvement (pN).

| experiment | count of cells before QC | | | | count of cells after basic QC | count of cells after filtering | metrics for CNA | | |
|---|---|---|---|---|---|---|---|---|---|
| | either assay | CNA only | ATAC only | both assay | both assay | both assay | average reads number | average PCR duplicate rate | average median bin count |
| wellDA-1 | 2,080 | 2,040 | 2,021 | 1,981 | 1,623 | 1,382 | 582,653 | 18.6% | 36 |
| wellDA-2 | 1,962 | 1,919 | 1,875 | 1,832 | 1,455 | 1,221 | 606,684 | 18.9% | 37 |

| experiment | metrics for ATAC | | | | | | |
|---|---|---|---|---|---|---|---|
| | average reads number | average number of fragments | average TSS enrichment score | average FRiP | average mono-nucleosomal fragments | average nucleosome-free fragments | overall PCR duplicate rate |
| wellDA-1 | 23,412 | 17,080 | 16.6 | 63.7% | 9,950 | 5,753 | 34.4% |
| wellDA-2 | 28,757 | 19,552 | 17.5 | 67.5% | 10,788 | 7,312 | 40.0% |

**Supplementary Table 2 QC metrics for MDA231 cell line**

QC and sequencing metrics of DNA and ATAC data from the two wellDA-seq experiments (wellDA-1 and wellDA-2) from the MDA231 cell line. ‘Before QC’: keeping cells that are found dispensed and have non-zero sequencing reads of wellDA-seq. ‘Basic QC’: keeping cells that simultaneously are found dispensed, passed the ATAC QC (>=100 fragments and >=8 TSS enrichment scores), and passed the CNA QC (>= 10 bin counts and having a correlation score > 0.8). ‘After filtering’: keeping cells that are not cell doublets or noise cells.

| Individual /patient | Sample_ID | count of cells passing QC | | | | metrics for CNA | | |
|---|---|---|---|---|---|---|---|---|
| | | both | either assay | CNA only | ATAC only | average reads number | average PCR duplicate rate | average median bin count |
| P1 | BCMHBCA52R | 1,339 | 1,525 | 1,465 | 1,489 | 821,436 | 20.3% | 50 |
| P2 | BCMHBCA66L | 1,115 | 1,398 | 1,308 | 1,221 | 316,782 | 16.4% | 20 |

| Individual /patient | metrics for ATAC | | | | | | |
|---|---|---|---|---|---|---|---|
| | average reads number | average number of fragments | average TSS enrichment score | average FRiP | average mono-nucleosomal fragments | average nucleosome-free fragments | overall PCR duplicate rate |
| P1 | 33,257 | 13,045 | 22.7 | 72.6% | 5,669 | 6,469 | 65.3% |
| P2 | 34,544 | 19,286 | 13.3 | 34.3% | 10,713 | 6,226 | 45.0% |

**Supplementary Table 3 QC metrics for 2 normal breast samples**

QC and sequencing metrics of DNA and ATAC data from wellDA-seq experiments of normal breast tissues (P1, P2) from two healthy women.

| Individual / patient | Sample ID | experiment | count of cells passing QC | | | | metrics for CNA | | |
|---|---|---|---|---|---|---|---|---|---|
| | | | both | either assay | CNA only | ATAC only | average reads number | average PCR duplicate rate | average median bin count |
| P3 | BCMDCIS22T | DCIS22T_chip1 | 1,477 | 2,062 | 2,035 | 1,773 | 555,003 | 12.1% | 37 |
| | | DCIS22T_chip2 | 1,392 | 2,126 | 2,093 | 1,822 | 559,080 | 17.7% | 35 |
| P4 | BCMDCIS28T | DCIS28T_chip1 | 915 | 1,212 | 1,203 | 1,006 | 949,890 | 17.1% | 60 |
| | | DCIS28T_chip2 | 956 | 1,443 | 1,427 | 1,089 | 826,639 | 21.1% | 49 |
| P5 | BCMDCI35T | DCIS35T_chip1 | 1,687 | 2,028 | 2,008 | 1,910 | 559,739 | 20.0% | 34 |
| | | DCIS35T_chip2 | 1,386 | 1,819 | 1,767 | 1,661 | 656,871 | 18.2% | 40 |
| P6 | BCMDCIS41T | DCIS41T_chip1 | 1,140 | 1,941 | 1,889 | 1,849 | 605,027 | 20.3% | 36 |
| | | DCIS41T_chip2 | 1,198 | 1,894 | 1,846 | 1,828 | 613,839 | 18.1% | 37 |
| P7 | BCMDCIS51T | DCIS51T_chip1 | 778 | 1,452 | 1,414 | 1,392 | 781,158 | 17.9% | 47 |
| | | DCIS51T_chip2 | 978 | 1,324 | 1,321 | 1,283 | 870,991 | 19.8% | 51 |
| P8 | BCMDCIS66T | DCIS66T_chip2 | 1,541 | 2,090 | 1,998 | 2,013 | 486,368 | 16.2% | 31 |
| P9 | BCMDCIS67T | DCIS67T_chip1 | 615 | 1,103 | 1,007 | 1,029 | 483,660 | 17.7% | 30 |
| | | DCIS67T_chip2 | 829 | 1,233 | 1,198 | 1,174 | 967,539 | 23.6% | 56 |
| P10 | BCMDCIS68T | DCIS68T_chip1 | 1,609 | 2,155 | 2,092 | 2,059 | 502,146 | 23.4% | 29 |
| | | DCIS68T_chip2 | 467 | 937 | 743 | 893 | 689,817 | 21.8% | 40 |
| P11 | BCMDCIS69T | BCMDCIS69_chip1 | 1,505 | 2,019 | 1,965 | 1,842 | 503,131 | 18.5% | 31 |
| | | BCMDCIS69_chip2 | 1,196 | 1,664 | 1,619 | 1,511 | 543,818 | 15.6% | 34 |

| Individual / patient | metrics for ATAC | | | | | | |
|---|---|---|---|---|---|---|---|
| | average reads number | average number of fragments | average TSS enrichment score | average FRiP | average mono-nucleosomal fragments | average nucleosome-free fragments | overall PCR duplicate rate |
| P3 | 13,881 | 9,515 | 16.8 | 32.3% | 5,461 | 3,306 | 34.4% |
| | 17,443 | 11,689 | 16.2 | 30.9% | 6,850 | 3,852 | 35.0% |
| P4 | 31,350 | 17,997 | 13.7 | 31.9% | 9,398 | 7,415 | 43.9% |
| | 22,874 | 15,718 | 11.6 | 26.9% | 8,952 | 5,165 | 32.7% |
| P5 | 18,321 | 9,518 | 18.4 | 50.9% | 3,611 | 5,234 | 50.7% |
| | 22,998 | 11,436 | 16.9 | 47.8% | 4,723 | 5,575 | 51.9% |
| P6 | 19,158 | 11,784 | 15.7 | 46.3% | 6,036 | 4,536 | 41.8% |
| | 20,063 | 11,910 | 16.2 | 48.1% | 5,808 | 5,212 | 43.7% |
| P7 | 22,072 | 12,003 | 18.3 | 46.1% | 6,506 | 4,518 | 48.5% |
| | 17,591 | 9,529 | 19.9 | 46.9% | 5,097 | 3,758 | 48.4% |
| P8 | 15,148 | 7,183 | 18.0 | 44.1% | 2,901 | 3,642 | 55.7% |
| P9 | 30,490 | 10,995 | 14.9 | 34.1% | 6,137 | 3,646 | 65.0% |
| | 36,052 | 14,580 | 15.7 | 39.5% | 7,929 | 5,144 | 60.6% |
| P10 | 16,018 | 7,851 | 18.3 | 49.2% | 4,445 | 2,616 | 52.2% |
| | 56,846 | 14,398 | 17.4 | 46.7% | 8,324 | 4,194 | 74.1% |
| P11 | 22,784 | 7,838 | 17.6 | 32.5% | 3,371 | 3,618 | 66.8% |
| | 21,785 | 7,094 | 16.0 | 30.8% | 3,267 | 3,144 | 69.8% |

**Supplementary Table 4 QC metrics for 9 breast tumor samples**

QC and sequencing metrics of DNA and ATAC data from wellDA-seq experiments of 9 breast tumor samples (P3-P11).

| ATAC-CB1 primer sets (S5XX) | | |
|---|---|---|
| **Well Position** | **Name** | **Sequence** |
| A5 | wellDA_ATAC_S5_1 | AATGATACGGCGACCACCGAGATCTACACTGA**GTATCGGT**GCCTCCCTCGCGCCAT |
| B5 | wellDA_ATAC_S5_2 | AATGATACGGCGACCACCGAGATCTACACTGACCGAAGCAGCCTCCCTCGCGCCAT |
| C5 | wellDA_ATAC_S5_3 | AATGATACGGCGACCACCGAGATCTACACTGAATATTATCGCCTCCCTCGCGCCAT |
| D5 | wellDA_ATAC_S5_4 | AATGATACGGCGACCACCGAGATCTACACTGACCACACACGCCTCCCTCGCGCCAT |
| E5 | wellDA_ATAC_S5_5 | AATGATACGGCGACCACCGAGATCTACACTGATTCAATCCGCCTCCCTCGCGCCAT |
| F5 | wellDA_ATAC_S5_6 | AATGATACGGCGACCACCGAGATCTACACTGAGCCGGTAGGCCTCCCTCGCGCCAT |
| G5 | wellDA_ATAC_S5_7 | AATGATACGGCGACCACCGAGATCTACACTGACTTCAGTGGCCTCCCTCGCGCCAT |
| H5 | wellDA_ATAC_S5_8 | AATGATACGGCGACCACCGAGATCTACACTGAAGGCACTTGCCTCCCTCGCGCCAT |
| I5 | wellDA_ATAC_S5_9 | AATGATACGGCGACCACCGAGATCTACACTGATTCCGCGGGCCTCCCTCGCGCCAT |
| J5 | wellDA_ATAC_S5_10 | AATGATACGGCGACCACCGAGATCTACACTGAAAAGTCCGGCCTCCCTCGCGCCAT |
| K5 | wellDA_ATAC_S5_11 | AATGATACGGCGACCACCGAGATCTACACTGAGGCTACCCGCCTCCCTCGCGCCAT |
| L5 | wellDA_ATAC_S5_12 | AATGATACGGCGACCACCGAGATCTACACTGACAATGTAGGCCTCCCTCGCGCCAT |
| M5 | wellDA_ATAC_S5_13 | AATGATACGGCGACCACCGAGATCTACACTGACAACGGCAGCCTCCCTCGCGCCAT |
| N5 | wellDA_ATAC_S5_14 | AATGATACGGCGACCACCGAGATCTACACTGACTCCGAACGCCTCCCTCGCGCCAT |
| O5 | wellDA_ATAC_S5_15 | AATGATACGGCGACCACCGAGATCTACACTGAAACACCTAGCCTCCCTCGCGCCAT |
| P5 | wellDA_ATAC_S5_16 | AATGATACGGCGACCACCGAGATCTACACTGAGTGCCATTGCCTCCCTCGCGCCAT |
| A6 | wellDA_ATAC_S5_17 | AATGATACGGCGACCACCGAGATCTACACTGAGCGCTGCTGCCTCCCTCGCGCCAT |
| B6 | wellDA_ATAC_S5_18 | AATGATACGGCGACCACCGAGATCTACACTGAGGCGTCTAGCCTCCCTCGCGCCAT |
| C6 | wellDA_ATAC_S5_19 | AATGATACGGCGACCACCGAGATCTACACTGAGTAATACAGCCTCCCTCGCGCCAT |
| D6 | wellDA_ATAC_S5_20 | AATGATACGGCGACCACCGAGATCTACACTGAACTTGTTGGCCTCCCTCGCGCCAT |
| E6 | wellDA_ATAC_S5_21 | AATGATACGGCGACCACCGAGATCTACACTGAAGTCCGAGGCCTCCCTCGCGCCAT |
| F6 | wellDA_ATAC_S5_22 | AATGATACGGCGACCACCGAGATCTACACTGAATTACCGTGCCTCCCTCGCGCCAT |
| G6 | wellDA_ATAC_S5_23 | AATGATACGGCGACCACCGAGATCTACACTGACCCTTGGAGCCTCCCTCGCGCCAT |
| H6 | wellDA_ATAC_S5_24 | AATGATACGGCGACCACCGAGATCTACACTGAGCCCACGTGCCTCCCTCGCGCCAT |
| I6 | wellDA_ATAC_S5_25 | AATGATACGGCGACCACCGAGATCTACACTGATCAAGAGCGCCTCCCTCGCGCCAT |
| J6 | wellDA_ATAC_S5_26 | AATGATACGGCGACCACCGAGATCTACACTGAATCGAATGGCCTCCCTCGCGCCAT |
| K6 | wellDA_ATAC_S5_27 | AATGATACGGCGACCACCGAGATCTACACTGAGGTGAAGGGCCTCCCTCGCGCCAT |
| L6 | wellDA_ATAC_S5_28 | AATGATACGGCGACCACCGAGATCTACACTGAACTTAGAAGCCTCCCTCGCGCCAT |
| M6 | wellDA_ATAC_S5_29 | AATGATACGGCGACCACCGAGATCTACACTGACGATAGGGGCCTCCCTCGCGCCAT |
| N6 | wellDA_ATAC_S5_30 | AATGATACGGCGACCACCGAGATCTACACTGAAATCTACAGCCTCCCTCGCGCCAT |
| O6 | wellDA_ATAC_S5_31 | AATGATACGGCGACCACCGAGATCTACACTGATCTGGCGAGCCTCCCTCGCGCCAT |
| P6 | wellDA_ATAC_S5_32 | AATGATACGGCGACCACCGAGATCTACACTGAACGCGCAGGCCTCCCTCGCGCCAT |
| A7 | wellDA_ATAC_S5_33 | AATGATACGGCGACCACCGAGATCTACACTGAATCCAGGAGCCTCCCTCGCGCCAT |
| B7 | wellDA_ATAC_S5_34 | AATGATACGGCGACCACCGAGATCTACACTGATATTTGCGGCCTCCCTCGCGCCAT |
| C7 | wellDA_ATAC_S5_35 | AATGATACGGCGACCACCGAGATCTACACTGACGTAGACCGCCTCCCTCGCGCCAT |
| D7 | wellDA_ATAC_S5_36 | AATGATACGGCGACCACCGAGATCTACACTGACTTTAACAGCCTCCCTCGCGCCAT |
| E7 | wellDA_ATAC_S5_37 | AATGATACGGCGACCACCGAGATCTACACTGATTGCCTAAGCCTCCCTCGCGCCAT |
| F7 | wellDA_ATAC_S5_38 | AATGATACGGCGACCACCGAGATCTACACTGAGAGACGTGGCCTCCCTCGCGCCAT |
| G7 | wellDA_ATAC_S5_39 | AATGATACGGCGACCACCGAGATCTACACTGATAAGTTACGCCTCCCTCGCGCCAT |
| H7 | wellDA_ATAC_S5_40 | AATGATACGGCGACCACCGAGATCTACACTGACCACCGTTGCCTCCCTCGCGCCAT |
| I7 | wellDA_ATAC_S5_41 | AATGATACGGCGACCACCGAGATCTACACTGAGAACTCGAGCCTCCCTCGCGCCAT |
| J7 | wellDA_ATAC_S5_42 | AATGATACGGCGACCACCGAGATCTACACTGATCACATGGGCCTCCCTCGCGCCAT |
| K7 | wellDA_ATAC_S5_43 | AATGATACGGCGACCACCGAGATCTACACTGACGCCAGATGCCTCCCTCGCGCCAT |
| L7 | wellDA_ATAC_S5_44 | AATGATACGGCGACCACCGAGATCTACACTGACTAGAAGTGCCTCCCTCGCGCCAT |
| M7 | wellDA_ATAC_S5_45 | AATGATACGGCGACCACCGAGATCTACACTGAACCGGCCCGCCTCCCTCGCGCCAT |
| N7 | wellDA_ATAC_S5_46 | AATGATACGGCGACCACCGAGATCTACACTGACCAGTTTAGCCTCCCTCGCGCCAT |
| O7 | wellDA_ATAC_S5_47 | AATGATACGGCGACCACCGAGATCTACACTGACGTACCAAGCCTCCCTCGCGCCAT |
| P7 | wellDA_ATAC_S5_48 | AATGATACGGCGACCACCGAGATCTACACTGAATGGTCATGCCTCCCTCGCGCCAT |
| A8 | wellDA_ATAC_S5_49 | AATGATACGGCGACCACCGAGATCTACACTGAGATCATTCGCCTCCCTCGCGCCAT |
| B8 | wellDA_ATAC_S5_50 | AATGATACGGCGACCACCGAGATCTACACTGAGGCACGGAGCCTCCCTCGCGCCAT |
| C8 | wellDA_ATAC_S5_51 | AATGATACGGCGACCACCGAGATCTACACTGATGATTGTTGCCTCCCTCGCGCCAT |
| D8 | wellDA_ATAC_S5_52 | AATGATACGGCGACCACCGAGATCTACACTGACCTGCGGGGCCTCCCTCGCGCCAT |
| E8 | wellDA_ATAC_S5_53 | AATGATACGGCGACCACCGAGATCTACACTGAACGATAACGCCTCCCTCGCGCCAT |
| F8 | wellDA_ATAC_S5_54 | AATGATACGGCGACCACCGAGATCTACACTGAGCAGCCTCGCCTCCCTCGCGCCAT |
| G8 | wellDA_ATAC_S5_55 | AATGATACGGCGACCACCGAGATCTACACTGAGCAAATTTGCCTCCCTCGCGCCAT |
| H8 | wellDA_ATAC_S5_56 | AATGATACGGCGACCACCGAGATCTACACTGAGACGTGGCGCCTCCCTCGCGCCAT |
| I8 | wellDA_ATAC_S5_57 | AATGATACGGCGACCACCGAGATCTACACTGAGGGCTTGGGCCTCCCTCGCGCCAT |
| J8 | wellDA_ATAC_S5_58 | AATGATACGGCGACCACCGAGATCTACACTGAATTGGGTCGCCTCCCTCGCGCCAT |
| K8 | wellDA_ATAC_S5_59 | AATGATACGGCGACCACCGAGATCTACACTGAAGCCGTTCGCCTCCCTCGCGCCAT |
| L8 | wellDA_ATAC_S5_60 | AATGATACGGCGACCACCGAGATCTACACTGACGGACTGCGCCTCCCTCGCGCCAT |
| M8 | wellDA_ATAC_S5_61 | AATGATACGGCGACCACCGAGATCTACACTGAGATTCCCAGCCTCCCTCGCGCCAT |
| N8 | wellDA_ATAC_S5_62 | AATGATACGGCGACCACCGAGATCTACACTGAACATTCGTGCCTCCCTCGCGCCAT |
| O8 | wellDA_ATAC_S5_63 | AATGATACGGCGACCACCGAGATCTACACTGAGGTTCAATGCCTCCCTCGCGCCAT |
| P8 | wellDA_ATAC_S5_64 | AATGATACGGCGACCACCGAGATCTACACTGACAGCAACGGCCTCCCTCGCGCCAT |
| A9 | wellDA_ATAC_S5_65 | AATGATACGGCGACCACCGAGATCTACACTGACCTTATGTGCCTCCCTCGCGCCAT |
| B9 | wellDA_ATAC_S5_66 | AATGATACGGCGACCACCGAGATCTACACTGAAGGTTGCCGCCTCCCTCGCGCCAT |
| C9 | wellDA_ATAC_S5_67 | AATGATACGGCGACCACCGAGATCTACACTGATTCGTGCAGCCTCCCTCGCGCCAT |
| D9 | wellDA_ATAC_S5_68 | AATGATACGGCGACCACCGAGATCTACACTGATCCATCAAGCCTCCCTCGCGCCAT |
| A10 | wellDA_ATAC_S5_69 | AATGATACGGCGACCACCGAGATCTACACTGAGTGTATCGGCCTCCCTCGCGCCAT |
| B10 | wellDA_ATAC_S5_70 | AATGATACGGCGACCACCGAGATCTACACTGATAACCAAGGCCTCCCTCGCGCCAT |
| C10 | wellDA_ATAC_S5_71 | AATGATACGGCGACCACCGAGATCTACACTGATTTCTAGCGCCTCCCTCGCGCCAT |
| D10 | wellDA_ATAC_S5_72 | AATGATACGGCGACCACCGAGATCTACACTGA**GTCGGACT**GCCTCCCTCGCGCCAT |

| ATAC-CB2 primer sets (N7XX) | | |
|---|---|---|
| **Well Position** | **Name** | **Sequence** |
| A13 | wellDA_ATAC_N7_1 | CAAGCAGAAGACGGCATACGAGAT**TCGCCTTA**CTTGCCAGCCCGCTCAG |
| B13 | wellDA_ATAC_N7_2 | CAAGCAGAAGACGGCATACGAGATCTAGTACGCTTGCCAGCCCGCTCAG |
| C13 | wellDA_ATAC_N7_3 | CAAGCAGAAGACGGCATACGAGATTTCTGCCTCTTGCCAGCCCGCTCAG |
| D13 | wellDA_ATAC_N7_4 | CAAGCAGAAGACGGCATACGAGATGCTCAGGACTTGCCAGCCCGCTCAG |
| E13 | wellDA_ATAC_N7_5 | CAAGCAGAAGACGGCATACGAGATAGGAGTCCCTTGCCAGCCCGCTCAG |
| F13 | wellDA_ATAC_N7_6 | CAAGCAGAAGACGGCATACGAGATCATGCCTACTTGCCAGCCCGCTCAG |
| G13 | wellDA_ATAC_N7_7 | CAAGCAGAAGACGGCATACGAGATGTAGAGAGCTTGCCAGCCCGCTCAG |
| H13 | wellDA_ATAC_N7_8 | CAAGCAGAAGACGGCATACGAGATCCTCTCTGCTTGCCAGCCCGCTCAG |
| I13 | wellDA_ATAC_N7_9 | CAAGCAGAAGACGGCATACGAGATAGCGTAGCCTTGCCAGCCCGCTCAG |
| J13 | wellDA_ATAC_N7_10 | CAAGCAGAAGACGGCATACGAGATCAGCCTCGCTTGCCAGCCCGCTCAG |
| K13 | wellDA_ATAC_N7_11 | CAAGCAGAAGACGGCATACGAGATTGCCTCTTCTTGCCAGCCCGCTCAG |
| L13 | wellDA_ATAC_N7_12 | CAAGCAGAAGACGGCATACGAGATTCCTCTACCTTGCCAGCCCGCTCAG |
| M13 | wellDA_ATAC_N7_13 | CAAGCAGAAGACGGCATACGAGATTCATGAGCCTTGCCAGCCCGCTCAG |
| N13 | wellDA_ATAC_N7_14 | CAAGCAGAAGACGGCATACGAGATCCTGAGATCTTGCCAGCCCGCTCAG |
| O13 | wellDA_ATAC_N7_15 | CAAGCAGAAGACGGCATACGAGATTAGCGAGTCTTGCCAGCCCGCTCAG |
| P13 | wellDA_ATAC_N7_16 | CAAGCAGAAGACGGCATACGAGATGTAGCTCCCTTGCCAGCCCGCTCAG |
| A14 | wellDA_ATAC_N7_17 | CAAGCAGAAGACGGCATACGAGATTACTACGCCTTGCCAGCCCGCTCAG |
| B14 | wellDA_ATAC_N7_18 | CAAGCAGAAGACGGCATACGAGATAGGCTCCGCTTGCCAGCCCGCTCAG |
| C14 | wellDA_ATAC_N7_19 | CAAGCAGAAGACGGCATACGAGATGCAGCGTACTTGCCAGCCCGCTCAG |
| D14 | wellDA_ATAC_N7_20 | CAAGCAGAAGACGGCATACGAGATCTGCGCATCTTGCCAGCCCGCTCAG |
| E14 | wellDA_ATAC_N7_21 | CAAGCAGAAGACGGCATACGAGATGAGCGCTACTTGCCAGCCCGCTCAG |
| F14 | wellDA_ATAC_N7_22 | CAAGCAGAAGACGGCATACGAGATCGCTCAGTCTTGCCAGCCCGCTCAG |
| G14 | wellDA_ATAC_N7_23 | CAAGCAGAAGACGGCATACGAGATGTCTTAGGCTTGCCAGCCCGCTCAG |
| H14 | wellDA_ATAC_N7_24 | CAAGCAGAAGACGGCATACGAGATACTGATCGCTTGCCAGCCCGCTCAG |
| I14 | wellDA_ATAC_N7_25 | CAAGCAGAAGACGGCATACGAGATTAGCTGCACTTGCCAGCCCGCTCAG |
| J14 | wellDA_ATAC_N7_26 | CAAGCAGAAGACGGCATACGAGATGACGTCGACTTGCCAGCCCGCTCAG |
| K14 | wellDA_ATAC_N7_27 | CAAGCAGAAGACGGCATACGAGATCTTAGCAACTTGCCAGCCCGCTCAG |
| L14 | wellDA_ATAC_N7_28 | CAAGCAGAAGACGGCATACGAGATTAACTTATCTTGCCAGCCCGCTCAG |
| M14 | wellDA_ATAC_N7_29 | CAAGCAGAAGACGGCATACGAGATCGAGTGATCTTGCCAGCCCGCTCAG |
| N14 | wellDA_ATAC_N7_30 | CAAGCAGAAGACGGCATACGAGATCTGTTAACCTTGCCAGCCCGCTCAG |
| O14 | wellDA_ATAC_N7_31 | CAAGCAGAAGACGGCATACGAGATCTACCATTCTTGCCAGCCCGCTCAG |
| P14 | wellDA_ATAC_N7_32 | CAAGCAGAAGACGGCATACGAGATACGTGCTCCTTGCCAGCCCGCTCAG |
| A15 | wellDA_ATAC_N7_33 | CAAGCAGAAGACGGCATACGAGATTGACGAAACTTGCCAGCCCGCTCAG |
| B15 | wellDA_ATAC_N7_34 | CAAGCAGAAGACGGCATACGAGATAATTCTTGCTTGCCAGCCCGCTCAG |
| C15 | wellDA_ATAC_N7_35 | CAAGCAGAAGACGGCATACGAGATGGCATTTCCTTGCCAGCCCGCTCAG |
| D15 | wellDA_ATAC_N7_36 | CAAGCAGAAGACGGCATACGAGATCTGCGAGGCTTGCCAGCCCGCTCAG |
| E15 | wellDA_ATAC_N7_37 | CAAGCAGAAGACGGCATACGAGATGAGGTGTACTTGCCAGCCCGCTCAG |
| F15 | wellDA_ATAC_N7_38 | CAAGCAGAAGACGGCATACGAGATAAATGACCCTTGCCAGCCCGCTCAG |
| G15 | wellDA_ATAC_N7_39 | CAAGCAGAAGACGGCATACGAGATTAAGATTGCTTGCCAGCCCGCTCAG |
| H15 | wellDA_ATAC_N7_40 | CAAGCAGAAGACGGCATACGAGATAAGGCACACTTGCCAGCCCGCTCAG |
| I15 | wellDA_ATAC_N7_41 | CAAGCAGAAGACGGCATACGAGATTAATAAGACTTGCCAGCCCGCTCAG |
| J15 | wellDA_ATAC_N7_42 | CAAGCAGAAGACGGCATACGAGATACTAAGTCCTTGCCAGCCCGCTCAG |
| K15 | wellDA_ATAC_N7_43 | CAAGCAGAAGACGGCATACGAGATGCTGGTCTCTTGCCAGCCCGCTCAG |
| L15 | wellDA_ATAC_N7_44 | CAAGCAGAAGACGGCATACGAGATCTGTATTTCTTGCCAGCCCGCTCAG |
| M15 | wellDA_ATAC_N7_45 | CAAGCAGAAGACGGCATACGAGATTTTCATAACTTGCCAGCCCGCTCAG |
| N15 | wellDA_ATAC_N7_46 | CAAGCAGAAGACGGCATACGAGATGACCCAAGCTTGCCAGCCCGCTCAG |
| O15 | wellDA_ATAC_N7_47 | CAAGCAGAAGACGGCATACGAGATTTATTTGGCTTGCCAGCCCGCTCAG |
| P15 | wellDA_ATAC_N7_48 | CAAGCAGAAGACGGCATACGAGATTTTAACGCCTTGCCAGCCCGCTCAG |
| A16 | wellDA_ATAC_N7_49 | CAAGCAGAAGACGGCATACGAGATCTTACTCCCTTGCCAGCCCGCTCAG |
| B16 | wellDA_ATAC_N7_50 | CAAGCAGAAGACGGCATACGAGATGGGAACCGCTTGCCAGCCCGCTCAG |
| C16 | wellDA_ATAC_N7_51 | CAAGCAGAAGACGGCATACGAGATCCCATGAGCTTGCCAGCCCGCTCAG |
| D16 | wellDA_ATAC_N7_52 | CAAGCAGAAGACGGCATACGAGATGCATTAAACTTGCCAGCCCGCTCAG |
| E16 | wellDA_ATAC_N7_53 | CAAGCAGAAGACGGCATACGAGATGACCGTTTCTTGCCAGCCCGCTCAG |
| F16 | wellDA_ATAC_N7_54 | CAAGCAGAAGACGGCATACGAGATTTTGGATCCTTGCCAGCCCGCTCAG |
| G16 | wellDA_ATAC_N7_55 | CAAGCAGAAGACGGCATACGAGATATCATCATCTTGCCAGCCCGCTCAG |
| H16 | wellDA_ATAC_N7_56 | CAAGCAGAAGACGGCATACGAGATCGTGTTGGCTTGCCAGCCCGCTCAG |
| I16 | wellDA_ATAC_N7_57 | CAAGCAGAAGACGGCATACGAGATTGTTGTTACTTGCCAGCCCGCTCAG |
| J16 | wellDA_ATAC_N7_58 | CAAGCAGAAGACGGCATACGAGATGGTTTACCCTTGCCAGCCCGCTCAG |
| K16 | wellDA_ATAC_N7_59 | CAAGCAGAAGACGGCATACGAGATGGTCGATGCTTGCCAGCCCGCTCAG |
| L16 | wellDA_ATAC_N7_60 | CAAGCAGAAGACGGCATACGAGATGTTCCCATCTTGCCAGCCCGCTCAG |
| M16 | wellDA_ATAC_N7_61 | CAAGCAGAAGACGGCATACGAGATATTGGCCGCTTGCCAGCCCGCTCAG |
| N16 | wellDA_ATAC_N7_62 | CAAGCAGAAGACGGCATACGAGATTCATTCCCCTTGCCAGCCCGCTCAG |
| O16 | wellDA_ATAC_N7_63 | CAAGCAGAAGACGGCATACGAGATCCGAATACCTTGCCAGCCCGCTCAG |
| P16 | wellDA_ATAC_N7_64 | CAAGCAGAAGACGGCATACGAGATATAGCTGACTTGCCAGCCCGCTCAG |
| A17 | wellDA_ATAC_N7_65 | CAAGCAGAAGACGGCATACGAGATAGATAAATCTTGCCAGCCCGCTCAG |
| B17 | wellDA_ATAC_N7_66 | CAAGCAGAAGACGGCATACGAGATATCTCGGGCTTGCCAGCCCGCTCAG |
| C17 | wellDA_ATAC_N7_67 | CAAGCAGAAGACGGCATACGAGATAGACATTACTTGCCAGCCCGCTCAG |
| D17 | wellDA_ATAC_N7_68 | CAAGCAGAAGACGGCATACGAGATGAATTGGCCTTGCCAGCCCGCTCAG |
| A18 | wellDA_ATAC_N7_69 | CAAGCAGAAGACGGCATACGAGATGCACGGCGCTTGCCAGCCCGCTCAG |
| B18 | wellDA_ATAC_N7_70 | CAAGCAGAAGACGGCATACGAGATTCGGTCAGCTTGCCAGCCCGCTCAG |
| C18 | wellDA_ATAC_N7_71 | CAAGCAGAAGACGGCATACGAGATTCGAAATGCTTGCCAGCCCGCTCAG |
| D18 | wellDA_ATAC_N7_72 | CAAGCAGAAGACGGCATACGAGAT**TGGCAAGC**CTTGCCAGCCCGCTCAG |

**Supplementary Table 5 wellDA-seq ATAC cell barcode sequences**

The well position (in the source plate), names and sequences of the first (ATAC-CB1) and second (ATAC-CB2) ATAC cell barcode primers. The nucleotides highlighted in bold indicate the ATAC cell barcode index sequence for exemplary primers.

| DNA-CB1 primer sets (S5XX) | | |
|---|---|---|
| **Well Position** | **Name** | **Sequence** |
| A5 | wellDA_DNA_S5_1 | AATGATACGGCGACCACCGAGATCTACAC**TAGATCGC**TCGTCGGCAGCGTC |
| B5 | wellDA_DNA_S5_2 | AATGATACGGCGACCACCGAGATCTACACTATCCTCTTCGTCGGCAGCGTC |
| C5 | wellDA_DNA_S5_3 | AATGATACGGCGACCACCGAGATCTACACAGAGTAGATCGTCGGCAGCGTC |
| D5 | wellDA_DNA_S5_4 | AATGATACGGCGACCACCGAGATCTACACGTAAGGAGTCGTCGGCAGCGTC |
| E5 | wellDA_DNA_S5_5 | AATGATACGGCGACCACCGAGATCTACACACTGCATATCGTCGGCAGCGTC |
| F5 | wellDA_DNA_S5_6 | AATGATACGGCGACCACCGAGATCTACACAAGGAGTATCGTCGGCAGCGTC |
| G5 | wellDA_DNA_S5_7 | AATGATACGGCGACCACCGAGATCTACACCTAAGCCTTCGTCGGCAGCGTC |
| H5 | wellDA_DNA_S5_8 | AATGATACGGCGACCACCGAGATCTACACCGTCTAATTCGTCGGCAGCGTC |
| I5 | wellDA_DNA_S5_9 | AATGATACGGCGACCACCGAGATCTACACTCTCTCCGTCGTCGGCAGCGTC |
| J5 | wellDA_DNA_S5_10 | AATGATACGGCGACCACCGAGATCTACACTCGACTAGTCGTCGGCAGCGTC |
| K5 | wellDA_DNA_S5_11 | AATGATACGGCGACCACCGAGATCTACACTTCTAGCTTCGTCGGCAGCGTC |
| L5 | wellDA_DNA_S5_12 | AATGATACGGCGACCACCGAGATCTACACCCTAGAGTTCGTCGGCAGCGTC |
| M5 | wellDA_DNA_S5_13 | AATGATACGGCGACCACCGAGATCTACACGCGTAAGATCGTCGGCAGCGTC |
| N5 | wellDA_DNA_S5_14 | AATGATACGGCGACCACCGAGATCTACACCTATTAAGTCGTCGGCAGCGTC |
| O5 | wellDA_DNA_S5_15 | AATGATACGGCGACCACCGAGATCTACACAAGGCTATTCGTCGGCAGCGTC |
| P5 | wellDA_DNA_S5_16 | AATGATACGGCGACCACCGAGATCTACACGAGCCTTATCGTCGGCAGCGTC |
| A6 | wellDA_DNA_S5_17 | AATGATACGGCGACCACCGAGATCTACACTTATGCGATCGTCGGCAGCGTC |
| B6 | wellDA_DNA_S5_18 | AATGATACGGCGACCACCGAGATCTACACTAAGCGTTTCGTCGGCAGCGTC |
| C6 | wellDA_DNA_S5_19 | AATGATACGGCGACCACCGAGATCTACACTCCGTCTTTCGTCGGCAGCGTC |
| D6 | wellDA_DNA_S5_20 | AATGATACGGCGACCACCGAGATCTACACTTCTGTGTTCGTCGGCAGCGTC |
| E6 | wellDA_DNA_S5_21 | AATGATACGGCGACCACCGAGATCTACACTCTGCTGTTCGTCGGCAGCGTC |
| F6 | wellDA_DNA_S5_22 | AATGATACGGCGACCACCGAGATCTACACTTGGAGGTTCGTCGGCAGCGTC |
| G6 | wellDA_DNA_S5_23 | AATGATACGGCGACCACCGAGATCTACACTGATACGTTCGTCGGCAGCGTC |
| H6 | wellDA_DNA_S5_24 | AATGATACGGCGACCACCGAGATCTACACTGCATAGTTCGTCGGCAGCGTC |
| I6 | wellDA_DNA_S5_25 | AATGATACGGCGACCACCGAGATCTACACTGCGATCTTCGTCGGCAGCGTC |
| J6 | wellDA_DNA_S5_26 | AATGATACGGCGACCACCGAGATCTACACTTCCTGCTTCGTCGGCAGCGTC |
| K6 | wellDA_DNA_S5_27 | AATGATACGGCGACCACCGAGATCTACACTAGTGACTTCGTCGGCAGCGTC |
| L6 | wellDA_DNA_S5_28 | AATGATACGGCGACCACCGAGATCTACACTACAGGATTCGTCGGCAGCGTC |
| M6 | wellDA_DNA_S5_29 | AATGATACGGCGACCACCGAGATCTACACTGTGGTTGTCGTCGGCAGCGTC |
| N6 | wellDA_DNA_S5_30 | AATGATACGGCGACCACCGAGATCTACACTACTAGTCTCGTCGGCAGCGTC |
| O6 | wellDA_DNA_S5_31 | AATGATACGGCGACCACCGAGATCTACACTCGAAGTGTCGTCGGCAGCGTC |
| P6 | wellDA_DNA_S5_32 | AATGATACGGCGACCACCGAGATCTACACTAACGCTGTCGTCGGCAGCGTC |
| A7 | wellDA_DNA_S5_33 | AATGATACGGCGACCACCGAGATCTACACTTGGTATGTCGTCGGCAGCGTC |
| B7 | wellDA_DNA_S5_34 | AATGATACGGCGACCACCGAGATCTACACTGAACTGGTCGTCGGCAGCGTC |
| C7 | wellDA_DNA_S5_35 | AATGATACGGCGACCACCGAGATCTACACTACTTCGGTCGTCGGCAGCGTC |
| D7 | wellDA_DNA_S5_36 | AATGATACGGCGACCACCGAGATCTACACTCTCACGGTCGTCGGCAGCGTC |
| E7 | wellDA_DNA_S5_37 | AATGATACGGCGACCACCGAGATCTACACGAGACGTGTCGTCGGCAGCGTC |
| F7 | wellDA_DNA_S5_38 | AATGATACGGCGACCACCGAGATCTACACTTGCCTAATCGTCGGCAGCGTC |
| G7 | wellDA_DNA_S5_39 | AATGATACGGCGACCACCGAGATCTACACCTTTAACATCGTCGGCAGCGTC |
| H7 | wellDA_DNA_S5_40 | AATGATACGGCGACCACCGAGATCTACACCGTAGACCTCGTCGGCAGCGTC |
| I7 | wellDA_DNA_S5_41 | AATGATACGGCGACCACCGAGATCTACACTATTTGCGTCGTCGGCAGCGTC |
| J7 | wellDA_DNA_S5_42 | AATGATACGGCGACCACCGAGATCTACACATCCAGGATCGTCGGCAGCGTC |
| K7 | wellDA_DNA_S5_43 | AATGATACGGCGACCACCGAGATCTACACTCTGGCGATCGTCGGCAGCGTC |
| L7 | wellDA_DNA_S5_44 | AATGATACGGCGACCACCGAGATCTACACAATCTACATCGTCGGCAGCGTC |
| M7 | wellDA_DNA_S5_45 | AATGATACGGCGACCACCGAGATCTACACCGATAGGGTCGTCGGCAGCGTC |
| N7 | wellDA_DNA_S5_46 | AATGATACGGCGACCACCGAGATCTACACGGTGAAGGTCGTCGGCAGCGTC |
| O7 | wellDA_DNA_S5_47 | AATGATACGGCGACCACCGAGATCTACACATCGAATGTCGTCGGCAGCGTC |
| P7 | wellDA_DNA_S5_48 | AATGATACGGCGACCACCGAGATCTACACTCAAGAGCTCGTCGGCAGCGTC |
| A8 | wellDA_DNA_S5_49 | AATGATACGGCGACCACCGAGATCTACACGCCCACGTTCGTCGGCAGCGTC |
| B8 | wellDA_DNA_S5_50 | AATGATACGGCGACCACCGAGATCTACACCCCTTGGATCGTCGGCAGCGTC |
| C8 | wellDA_DNA_S5_51 | AATGATACGGCGACCACCGAGATCTACACATTACCGTTCGTCGGCAGCGTC |
| D8 | wellDA_DNA_S5_52 | AATGATACGGCGACCACCGAGATCTACACAGTCCGAGTCGTCGGCAGCGTC |
| E8 | wellDA_DNA_S5_53 | AATGATACGGCGACCACCGAGATCTACACACTTGTTGTCGTCGGCAGCGTC |
| F8 | wellDA_DNA_S5_54 | AATGATACGGCGACCACCGAGATCTACACGTAATACATCGTCGGCAGCGTC |

| | | |
|---|---|---|
| G8 | wellDA_DNA_S5_55 | AATGATACGGCGACCACCGAGATCTACACGGCGTCTATCGTCGGCAGCGTC |
| H8 | wellDA_DNA_S5_56 | AATGATACGGCGACCACCGAGATCTACACGCGCTGCTTCGTCGGCAGCGTC |
| I8 | wellDA_DNA_S5_57 | AATGATACGGCGACCACCGAGATCTACACGTGCCATTTCGTCGGCAGCGTC |
| J8 | wellDA_DNA_S5_58 | AATGATACGGCGACCACCGAGATCTACACAACACCTATCGTCGGCAGCGTC |
| K8 | wellDA_DNA_S5_59 | AATGATACGGCGACCACCGAGATCTACACCTCCGAACTCGTCGGCAGCGTC |
| L8 | wellDA_DNA_S5_60 | AATGATACGGCGACCACCGAGATCTACACCAACGGCATCGTCGGCAGCGTC |
| M8 | wellDA_DNA_S5_61 | AATGATACGGCGACCACCGAGATCTACACCAATGTAGTCGTCGGCAGCGTC |
| N8 | wellDA_DNA_S5_62 | AATGATACGGCGACCACCGAGATCTACACGGCTACCCTCGTCGGCAGCGTC |
| O8 | wellDA_DNA_S5_63 | AATGATACGGCGACCACCGAGATCTACACAAAGTCCGTCGTCGGCAGCGTC |
| P8 | wellDA_DNA_S5_64 | AATGATACGGCGACCACCGAGATCTACACTTCCGCGGTCGTCGGCAGCGTC |
| A9 | wellDA_DNA_S5_65 | AATGATACGGCGACCACCGAGATCTACACAGGCACTTTCGTCGGCAGCGTC |
| B9 | wellDA_DNA_S5_66 | AATGATACGGCGACCACCGAGATCTACACCTTCAGTGTCGTCGGCAGCGTC |
| C9 | wellDA_DNA_S5_67 | AATGATACGGCGACCACCGAGATCTACACGCCGGTAGTCGTCGGCAGCGTC |
| D9 | wellDA_DNA_S5_68 | AATGATACGGCGACCACCGAGATCTACACTTCAATCCTCGTCGGCAGCGTC |
| A10 | wellDA_DNA_S5_69 | AATGATACGGCGACCACCGAGATCTACACCCACACACTCGTCGGCAGCGTC |
| B10 | wellDA_DNA_S5_70 | AATGATACGGCGACCACCGAGATCTACACATATTATCTCGTCGGCAGCGTC |
| C10 | wellDA_DNA_S5_71 | AATGATACGGCGACCACCGAGATCTACACCCGAAGCATCGTCGGCAGCGTC |
| D10 | wellDA_DNA_S5_72 | AATGATACGGCGACCACCGAGATCTACAC**GTATCGGT**TCGTCGGCAGCGTC |

### DNA-CB2 primer sets (N7XX)

| **Well Position** | **Name** | **Sequence** |
|---|---|---|
| A13 | wellDA_DNA_IN_N7_1 | CTGAGTCGGAGACACGCA**ACAAACGG**GTCTCGTGGGCTCGG |
| B13 | wellDA_DNA_IN_N7_2 | CTGAGTCGGAGACACGCAACCCAGCAGTCTCGTGGGCTCGG |
| C13 | wellDA_DNA_IN_N7_3 | CTGAGTCGGAGACACGCACCCAACCTGTCTCGTGGGCTCGG |
| D13 | wellDA_DNA_IN_N7_4 | CTGAGTCGGAGACACGCACACCACACGTCTCGTGGGCTCGG |
| E13 | wellDA_DNA_IN_N7_5 | CTGAGTCGGAGACACGCAGAAACCCAGTCTCGTGGGCTCGG |
| F13 | wellDA_DNA_IN_N7_6 | CTGAGTCGGAGACACGCATGTGACCAGTCTCGTGGGCTCGG |
| G13 | wellDA_DNA_IN_N7_7 | CTGAGTCGGAGACACGCAAGGGTCAAGTCTCGTGGGCTCGG |
| H13 | wellDA_DNA_IN_N7_8 | CTGAGTCGGAGACACGCATGTCCGCGGTCTCGTGGGCTCGG |
| I13 | wellDA_DNA_IN_N7_9 | CTGAGTCGGAGACACGCAATATGGAAGTCTCGTGGGCTCGG |
| J13 | wellDA_DNA_IN_N7_10 | CTGAGTCGGAGACACGCAAACGAATTGTCTCGTGGGCTCGG |
| K13 | wellDA_DNA_IN_N7_11 | CTGAGTCGGAGACACGCATCGACGCCGTCTCGTGGGCTCGG |
| L13 | wellDA_DNA_IN_N7_12 | CTGAGTCGGAGACACGCACACTTTGTGTCTCGTGGGCTCGG |
| M13 | wellDA_DNA_IN_N7_13 | CTGAGTCGGAGACACGCATTCAAGTAGTCTCGTGGGCTCGG |
| N13 | wellDA_DNA_IN_N7_14 | CTGAGTCGGAGACACGCAGCTATCACGTCTCGTGGGCTCGG |
| O13 | wellDA_DNA_IN_N7_15 | CTGAGTCGGAGACACGCAAATCTACTGTCTCGTGGGCTCGG |
| P13 | wellDA_DNA_IN_N7_16 | CTGAGTCGGAGACACGCACCGGCAATGTCTCGTGGGCTCGG |
| A14 | wellDA_DNA_IN_N7_17 | CTGAGTCGGAGACACGCACTTAGCAAGTCTCGTGGGCTCGG |
| B14 | wellDA_DNA_IN_N7_18 | CTGAGTCGGAGACACGCATAACTTATGTCTCGTGGGCTCGG |
| C14 | wellDA_DNA_IN_N7_19 | CTGAGTCGGAGACACGCACGAGTGATGTCTCGTGGGCTCGG |
| D14 | wellDA_DNA_IN_N7_20 | CTGAGTCGGAGACACGCACTGTTAACGTCTCGTGGGCTCGG |
| E14 | wellDA_DNA_IN_N7_21 | CTGAGTCGGAGACACGCACTACCATTGTCTCGTGGGCTCGG |
| F14 | wellDA_DNA_IN_N7_22 | CTGAGTCGGAGACACGCAACGTGCTCGTCTCGTGGGCTCGG |
| G14 | wellDA_DNA_IN_N7_23 | CTGAGTCGGAGACACGCATGACGAAAGTCTCGTGGGCTCGG |
| H14 | wellDA_DNA_IN_N7_24 | CTGAGTCGGAGACACGCAAATTCTTGGTCTCGTGGGCTCGG |
| I14 | wellDA_DNA_IN_N7_25 | CTGAGTCGGAGACACGCAGGCATTTCGTCTCGTGGGCTCGG |
| J14 | wellDA_DNA_IN_N7_26 | CTGAGTCGGAGACACGCACTGCGAGGGTCTCGTGGGCTCGG |
| K14 | wellDA_DNA_IN_N7_27 | CTGAGTCGGAGACACGCAGAGGTGTAGTCTCGTGGGCTCGG |
| L14 | wellDA_DNA_IN_N7_28 | CTGAGTCGGAGACACGCAAAATGACCGTCTCGTGGGCTCGG |
| M14 | wellDA_DNA_IN_N7_29 | CTGAGTCGGAGACACGCATAAGATTGGTCTCGTGGGCTCGG |
| N14 | wellDA_DNA_IN_N7_30 | CTGAGTCGGAGACACGCAAAGGCACAGTCTCGTGGGCTCGG |
| O14 | wellDA_DNA_IN_N7_31 | CTGAGTCGGAGACACGCATAATAAGAGTCTCGTGGGCTCGG |
| P14 | wellDA_DNA_IN_N7_32 | CTGAGTCGGAGACACGCAACTAAGTCGTCTCGTGGGCTCGG |
| A15 | wellDA_DNA_IN_N7_33 | CTGAGTCGGAGACACGCAGCTGGTCTGTCTCGTGGGCTCGG |
| B15 | wellDA_DNA_IN_N7_34 | CTGAGTCGGAGACACGCACTGTATTTGTCTCGTGGGCTCGG |
| C15 | wellDA_DNA_IN_N7_35 | CTGAGTCGGAGACACGCATTTCATAAGTCTCGTGGGCTCGG |
| D15 | wellDA_DNA_IN_N7_36 | CTGAGTCGGAGACACGCAGACCCAAGGTCTCGTGGGCTCGG |

| | | |
|---|---|---|
| E15 | wellDA_DNA_IN_N7_37 | CTGAGTCGGAGACACGCATTATTTGGGTCTCGTGGGCTCGG |
| F15 | wellDA_DNA_IN_N7_38 | CTGAGTCGGAGACACGCATTTAACGCGTCTCGTGGGCTCGG |
| G15 | wellDA_DNA_IN_N7_39 | CTGAGTCGGAGACACGCACTTACTCCGTCTCGTGGGCTCGG |
| H15 | wellDA_DNA_IN_N7_40 | CTGAGTCGGAGACACGCAGGGAACCGGTCTCGTGGGCTCGG |
| I15 | wellDA_DNA_IN_N7_41 | CTGAGTCGGAGACACGCACCCATGAGGTCTCGTGGGCTCGG |
| J15 | wellDA_DNA_IN_N7_42 | CTGAGTCGGAGACACGCAGCATTAAAGTCTCGTGGGCTCGG |
| K15 | wellDA_DNA_IN_N7_43 | CTGAGTCGGAGACACGCAGACCGTTTGTCTCGTGGGCTCGG |
| L15 | wellDA_DNA_IN_N7_44 | CTGAGTCGGAGACACGCATTTGGATCGTCTCGTGGGCTCGG |
| M15 | wellDA_DNA_IN_N7_45 | CTGAGTCGGAGACACGCAATCATCATGTCTCGTGGGCTCGG |
| N15 | wellDA_DNA_IN_N7_46 | CTGAGTCGGAGACACGCACGTGTTGGGTCTCGTGGGCTCGG |
| O15 | wellDA_DNA_IN_N7_47 | CTGAGTCGGAGACACGCATGTTGTTAGTCTCGTGGGCTCGG |
| P15 | wellDA_DNA_IN_N7_48 | CTGAGTCGGAGACACGCAGGTTTACCGTCTCGTGGGCTCGG |
| A16 | wellDA_DNA_IN_N7_49 | CTGAGTCGGAGACACGCAGGTCGATGGTCTCGTGGGCTCGG |
| B16 | wellDA_DNA_IN_N7_50 | CTGAGTCGGAGACACGCAGTTCCCATGTCTCGTGGGCTCGG |
| C16 | wellDA_DNA_IN_N7_51 | CTGAGTCGGAGACACGCAATTGGCCGGTCTCGTGGGCTCGG |
| D16 | wellDA_DNA_IN_N7_52 | CTGAGTCGGAGACACGCATCATTCCCGTCTCGTGGGCTCGG |
| E16 | wellDA_DNA_IN_N7_53 | CTGAGTCGGAGACACGCACCGAATACGTCTCGTGGGCTCGG |
| F16 | wellDA_DNA_IN_N7_54 | CTGAGTCGGAGACACGCAATAGCTGAGTCTCGTGGGCTCGG |
| G16 | wellDA_DNA_IN_N7_55 | CTGAGTCGGAGACACGCAAGATAAATGTCTCGTGGGCTCGG |
| H16 | wellDA_DNA_IN_N7_56 | CTGAGTCGGAGACACGCAATCTCGGGGTCTCGTGGGCTCGG |
| I16 | wellDA_DNA_IN_N7_57 | CTGAGTCGGAGACACGCAAGACATTAGTCTCGTGGGCTCGG |
| J16 | wellDA_DNA_IN_N7_58 | CTGAGTCGGAGACACGCAGAATTGGCGTCTCGTGGGCTCGG |
| K16 | wellDA_DNA_IN_N7_59 | CTGAGTCGGAGACACGCAGCACGGCGGTCTCGTGGGCTCGG |
| L16 | wellDA_DNA_IN_N7_60 | CTGAGTCGGAGACACGCATCGGTCAGGTCTCGTGGGCTCGG |
| M16 | wellDA_DNA_IN_N7_61 | CTGAGTCGGAGACACGCATCGAAATGGTCTCGTGGGCTCGG |
| N16 | wellDA_DNA_IN_N7_62 | CTGAGTCGGAGACACGCATGGCAAGCGTCTCGTGGGCTCGG |
| O16 | wellDA_DNA_IN_N7_63 | CTGAGTCGGAGACACGCATGGTAGAAGTCTCGTGGGCTCGG |
| P16 | wellDA_DNA_IN_N7_64 | CTGAGTCGGAGACACGCACGTCACGTGTCTCGTGGGCTCGG |
| A17 | wellDA_DNA_IN_N7_65 | CTGAGTCGGAGACACGCACGCGGACAGTCTCGTGGGCTCGG |
| B17 | wellDA_DNA_IN_N7_66 | CTGAGTCGGAGACACGCAAAGTTTAAGTCTCGTGGGCTCGG |
| C17 | wellDA_DNA_IN_N7_67 | CTGAGTCGGAGACACGCAGTTGTGGTGTCTCGTGGGCTCGG |
| D17 | wellDA_DNA_IN_N7_68 | CTGAGTCGGAGACACGCACCAGAGGCGTCTCGTGGGCTCGG |
| A18 | wellDA_DNA_IN_N7_69 | CTGAGTCGGAGACACGCAGTGGGCGAGTCTCGTGGGCTCGG |
| B18 | wellDA_DNA_IN_N7_70 | CTGAGTCGGAGACACGCAGCCTAGTGGTCTCGTGGGCTCGG |
| C18 | wellDA_DNA_IN_N7_71 | CTGAGTCGGAGACACGCAGTACATGCGTCTCGTGGGCTCGG |
| D18 | wellDA_DNA_IN_N7_72 | CTGAGTCGGAGACACGCA**ACTCGAAC**GTCTCGTGGGCTCGG |

| DNA enrichment PCR primer | | |
|---|---|---|
| | **Name** | **Sequence** |
| | wellDA_DNA_OUT_N7_1 | CAAGCAGAAGACGGCATACGAGAT**TCGCCTTA**CTGAGTCGGAGACACGCA |
| | wellDA_DNA_OUT_N7_2 | CAAGCAGAAGACGGCATACGAGATCTAGTACGCTGAGTCGGAGACACGCA |
| | wellDA_DNA_OUT_N7_3 | CAAGCAGAAGACGGCATACGAGATTTCTGCCTCTGAGTCGGAGACACGCA |
| | wellDA_DNA_OUT_N7_4 | CAAGCAGAAGACGGCATACGAGATGCTCAGGACTGAGTCGGAGACACGCA |
| | wellDA_DNA_OUT_N7_5 | CAAGCAGAAGACGGCATACGAGATAGGAGTCCCTGAGTCGGAGACACGCA |
| | wellDA_DNA_OUT_N7_6 | CAAGCAGAAGACGGCATACGAGATGTAGAGAGCTGAGTCGGAGACACGCA |
| | wellDA_DNA_OUT_N7_7 | CAAGCAGAAGACGGCATACGAGATCCTCTCTGCTGAGTCGGAGACACGCA |
| | wellDA_DNA_OUT_N7_8 | CAAGCAGAAGACGGCATACGAGATAGCGTAGCCTGAGTCGGAGACACGCA |
| | wellDA_DNA_OUT_N7_9 | CAAGCAGAAGACGGCATACGAGATCAGCCTCGCTGAGTCGGAGACACGCA |
| | wellDA_DNA_OUT_N7_10 | CAAGCAGAAGACGGCATACGAGATTGCCTCTTCTGAGTCGGAGACACGCA |
| | wellDA_DNA_OUT_N7_11 | CAAGCAGAAGACGGCATACGAGAT**TCCTCTAC**CTGAGTCGGAGACACGCA |

**Supplementary Table 6 wellDA-seq DNA cell barcode sequences**

The well position (in the source plate), names and sequences of the first (DNA-CB1), second (DNA-CB2) DNA cell barcode primers and the enrichment PCR primers. The nucleotides highlighted in bold indicate the DNA index sequence for exemplary primers.
